# A small molecule inhibitor of CD28 costimulation restrains pathogenic T-cell responses in inflammatory bowel disease

**DOI:** 10.64898/2026.08.10.744081

**Authors:** Sungwoo Cho, Saurabh Upadhyay, Shaoren Yuan, Moustafa Gabr

## Abstract

CD28 costimulation contributes to pathogenic T cell responses in inflammatory bowel disease (IBD), but current B7-directed blockade also limits CTLA-4 signaling. Using a sensitive NanoBiT split-luciferase screening platform, we identified and optimized CA-23, a small molecule antagonist that directly binds human and mouse CD28 without measurable binding to CD80, CD86, or CTLA-4. CA-23 inhibited CD28-B7 engagement and CD28-dependent T cell activation without agonist activity in human whole blood and peripheral blood mononuclear cells. CA-23 achieved exposure in the colon and mesenteric lymph nodes and reduced disease severity, histologic injury, and pathogenic Th1 and Th17 responses in a T cell transfer model of colitis. In PBMCs from donors with ulcerative colitis or Crohn’s disease, CA-23 suppressed inflammatory cytokine production and T cell activation to a degree matching or exceeding Abatacept. In human intestinal epithelial–PBMC co-cultures, CA-23 preserved Treg suppressive activity and epithelial barrier integrity, whereas Abatacept reduced Treg function. CA-23 did not alter CD80 or CD86 expression on autologous antigen-presenting cells and showed no substantial off-target activity in the tested selectivity panel. These findings support direct CD28 antagonism as a mechanistically differentiated alternative to B7-directed co-stimulation blockade for suppressing pathogenic T cell responses in preclinical models of IBD.

**One Sentence Summary:** A CD28-selective small molecule blocks pathogenic T cell activation and preserves Treg function unlike Abatacept in IBD models.

## INTRODUCTION

The activation of naïve T cells requires the integration of two distinct signals at the immunological synapse: antigen recognition through the T cell receptor (TCR), and a costimulatory signal delivered predominantly by CD28, a homodimeric type I transmembrane glycoprotein constitutively expressed on most resting T cells (*1*, *2*). CD28 engages two structurally homologous ligands of the B7 family, CD80 (B7-1) and CD86 (B7-2), which are upregulated on professional antigen-presenting cells (APCs) during innate immune activation (*3*, *4*). Genetic ablation of CD28 in mice abrogates productive T cell responses to a wide range of antigenic challenges (*5*), and TCR engagement in the absence of CD28 co-stimulation drives T cells into a state of clonal anergy rather than productive activation (*5*, *6*). At the molecular level, CD28 ligation amplifies TCR-induced PI3K, NF-κB, and NFAT signaling, reinforcing clonal expansion, IL-2 production, and effector differentiation (*7*, *8*). Consistent with this central role, the CD28-B7 axis is now recognized as a master regulator of pathogenic T cell responses underlying chronic inflammatory and autoimmune diseases.

Among such conditions, inflammatory bowel disease (IBD), encompassing Crohn’s disease and ulcerative colitis, exemplifies the clinical consequences of dysregulated T cell activation in mucosal tissues. IBD is a chronic, relapsing disorder of the gastrointestinal tract with a substantial and growing global health burden, driven in large part by aberrant expansion of pathogenic effector CD4+ T cell subsets within the intestinal lamina propria and gut-draining lymphoid tissues (*9*). Mouse colitis models and human IBD tissue have established central pathogenic roles for IFN-γ-producing Th1 cells, IL-17A-producing Th17 cells, and a particularly pathogenic IFN-γ+IL-17A+ double-producing CD4+ subset that emerges through Th17-to-Th1 plasticity at chronic mucosal sites (*10–14*). Although biologics that neutralize downstream cytokines or block lymphocyte trafficking, including anti-TNF-α, anti-IL-12/23, and anti-integrin antibodies, have transformed management of moderate-to-severe IBD, a substantial proportion of patients exhibit primary non-response, secondary loss of response, or treatment-limiting adverse events, leaving a clear unmet need for therapies that intervene further upstream in the T cell activation cascade (*15*, *16*).

Targeting CD28 co-stimulation directly, rather than its downstream cytokine effectors, represents a conceptually attractive upstream strategy. The only clinically approved co-stimulation blocker, Abatacept, is a soluble CTLA-4-Ig fusion protein that binds CD80 and CD86 on APCs and thereby prevents their engagement of CD28; it received FDA approval for rheumatoid arthritis in 2005 and is now also approved for juvenile idiopathic and psoriatic arthritis (*17–19*). Critically, however, Abatacept does not engage CD28 directly. The inhibitory coinhibitory receptor CTLA-4, which is constitutively expressed on regulatory T cells (Tregs) and is required for Treg-mediated immune suppression, shares CD80/CD86 as ligands and binds them with substantially higher affinity than CD28 itself (*20–22*). Consequently, Abatacept indiscriminately occupies B7 ligands and disrupts both stimulatory CD28 and inhibitory CTLA-4 engagement. Multiple clinical and mechanistic studies have correspondingly shown that Abatacept treatment reduces Treg frequency and impairs Treg suppressive function in patients, representing an intrinsic off-target liability of its B7-directed mechanism (*23*, *24*). A small molecule antagonist that selectively engages CD28, sparing the CTLA-4 axis and preserving Treg-mediated tolerance, would therefore represent a mechanistically differentiated and potentially safer modality. Despite this rationale, no small molecule CD28 antagonist has reached clinical development, in part because the CD28 -B7 interface, like many protein-protein interactions (PPIs), presents a flat, solvent-exposed contact area that lacks the deep ligand-binding pockets characteristic of conventional drug targets and has historically been considered “undruggable” (*25*, *26*).

Discovery of small molecule disruptors of such PPI interfaces requires screening platforms sensitive enough to detect partial inhibition of relatively weak and transient receptor-ligand contacts, while remaining compatible with high-throughput screening (HTS) workflows. The NanoBiT split-luciferase system addresses this by splitting NanoLuc into Large BiT (LgBiT) and Small BiT (SmBiT), whose weak intrinsic affinity (K_D_ ≈ 190 μM) permits reconstitution of luminescence only upon a genuine interaction between the fusion partners onto which they are appended (*27*). The same group later identified HiBiT, a higher-affinity SmBiT variant (K_D_ ≈ 700 pM) that substantially improves signal brightness, sensitivity, and dynamic range for PPI monitoring at near-physiological expression levels (*28*).

Here, we applied a HiBiT-LgBiT-based NanoBiT platform to perform, to our knowledge, the first HTS campaign for direct, CD28-selective small molecule inhibitors of the CD28-B7 axis. Screening of approximately 5,000 structurally diverse compounds identified two distinct sub-micromolar hit scaffolds, which we optimized through focused structure-activity relationship studies to nanomolar-potency lead compounds. Our final lead, CA-23, binds CD28 directly with sub-micromolar affinity and exhibits no detectable binding to CD80, CD86, or CTLA-4 even at 100 μM, thereby directly addressing the central selectivity limitation of Abatacept. CA-23 inhibits trans CD28-B7 engagement and downstream T cell activation across multiple orthogonal cellular and biochemical assays at sub-micromolar potency. In a CD4^+^CD45RB^hi^ T cell adoptive transfer model of chronic colitis, CA-23 dose-dependently attenuated weight loss, restored colon length, reduced histopathological inflammation, and suppressed the expansion of pathogenic Th1, Th17, and IFN-γ^+^IL-17A^+^ double-producing CD4^+^ T cells in the disease-proximal mesenteric lymph nodes. Collectively, these findings establish CA-23 as a novel CD28-selective small molecule antagonist with demonstrated efficacy in a T cell-driven in vivo model of IBD and provide a chemical and translational foundation for the development of next-generation, mechanism-differentiated CD28-directed immunomodulators.

## RESULTS

### A HiBiT-LgBiT NanoBiT assay for high-throughput screening of CD28-B7 interaction inhibitors

To enable high-throughput screening (HTS) of small molecule inhibitors targeting the CD28-B7 protein-protein interaction (PPI), we adapted the NanoBiT split-luciferase complementation system to monitor CD28-B7 engagement in a cell-based format. CD28 was fused to HiBiT, while the B7 ligands CD80 and CD86 were each fused to LgBiT, such that productive CD28-B7 engagement reconstitutes functional NanoLuc (Nanoluciferase) and generates luminescence at 460 nm, which is abolished upon disruption of the interaction by a small molecule antagonist (Fig. 1A, B). Whereas our previous CD28 antagonist study employed the SmBiT fragment, here we applied the HiBiT-LgBiT configuration to the CD28-B7 system, taking advantage of the markedly higher intrinsic affinity of HiBiT for LgBiT (K_D_ in the low nanomolar range compared with ∼190 μM for SmBiT). This adaptation yielded a substantially improved signal-to-background ratio and greater sensitivity for detecting weak or transient PPI disruption events, critical for HTS of chemically diverse small molecule libraries. The HiBiT-CD28 and LgBiT-CD80/LgBiT-CD86 constructs were expressed in CHO-K1 cells, and assay performance was validated using Abatacept (CTLA-4-Ig), a clinically approved CD28-B7 blocker. Abatacept (50 μg/ml) robustly suppressed luminescence in both CD28-CD80 and CD28-CD86 NanoBiT pairs (***p < 0.001), confirming that the assay faithfully reports CD28-B7 engagement (Fig. 1C), with Z’ factors of 0.756 (CD28-CD80) and 0.621 (CD28-CD86), both exceeding the conventional HTS threshold (Z’ > 0.5). Together, these results establish a sensitive, reliable HiBiT-LgBiT NanoBiT platform for HTS-based identification of CD28-B7 PPI inhibitors.

**Fig. 1.**
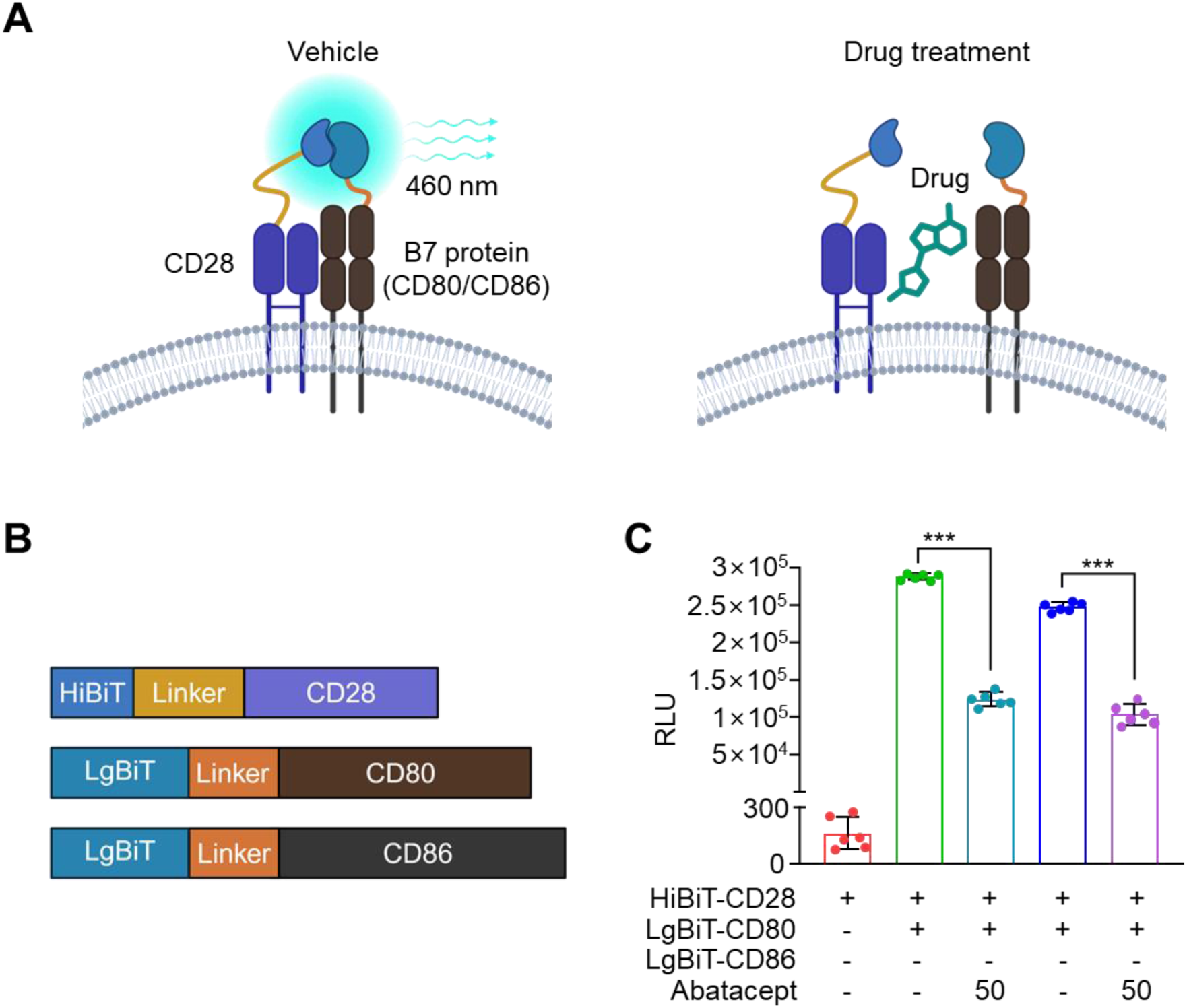
Development of a NanoBiT-based CD28-B7 interaction assay for compound screening. **(A)** Schematic representation of the NanoBiT assay. CD28 fused to HiBiT (small subunit) and B7 protein (CD80 or CD86) fused to LgBiT (large subunit) reconstitute functional Nano luciferase upon CD28-B7 interaction, producing luminescence at 460 nm (left). Drug treatment disrupts the CD28-B7 interaction, preventing NanoBiT reconstitution and abolishing luminescence (right). **(B)** Schematic of the fusion constructs: HiBiT-Linker-CD28, LgBiT-Linker-CD80, and LgBiT-Linker-CD86. **(C)** Validation of the assay using Abatacept (CTLA-4-Ig) at 50 μg/ml. CHOK-1 cells were co-transfected with the indicated constructs, and luminescence (RLU, relative luminescence units) was measured. Data are presented as mean ± SD (n=5). ***p < 0.001. analyzed by Student’s unpaired two-tailed t test.

### High-throughput screening identifies small molecule CD28-CD80 and CD28-CD86 inhibitors

To identify for CD28-B7 interaction inhibitor development, we screened the Enamine Discovery Diversity Set, a structurally diverse small molecule library, using the validated HiBiT-LgBiT based NanoBiT assay. For improved throughput, approximately 5,000 compounds were screened in a 4 - in-1 pooled format at a final concentration of 25 μM per compound against both the CD28-CD80 and CD28-CD86 NanoBiT pairs (Supplementary Fig. S1A, B). Primary hits were defined as exhibiting greater than 70% reduction in luminescence relative to vehicle (DMSO) controls, yielding 36 hits for CD28-CD80 and 65 hits for CD28-CD86. Counter-screening of the 25 hits overlapping between the CD28-CD80 and CD28-CD86 against our previously established NOS1-NOS1AP NanoBiT assay in CHO-K1 cells was performed to exclude nonspecific or assay-interfering compounds; 9 compounds that showed no activity against the unrelated NOS1 - NOS1AP PPI were retained as specific CD28-B7 inhibitor candidates (Supplementary Fig. S1C, Supplementary Fig. S2) (*29*). To further prioritize the most potent leads, all 9 candidates were re-tested at 10 μM in deconvoluted single-compound format, which identified CA-01 and CB-01 as the two compounds retaining strong inhibitory activity against both CD28-CD80 and CD28-CD86 interactions (Supplementary Fig. S1D-F). Full dose-response analysis confirmed potent and balanced inhibition of both interactions: CA-01 displayed IC50 values of 0.283 ± 0.03 μM (CD28-CD80) and 0.547 ± 0.21 μM (CD28-CD86) (Supplementary Fig. S1G), whereas CB-01 displayed IC_50_ values of 0.35 ± 0.16 μM (CD28-CD80) and 0.4 ± 0.06 μM (CD28-CD86) (Supplementary Fig. S1H). These two structurally distinct sub-micromolar inhibitors were thus selected as the primary hit scaffolds for subsequent structure-activity relationship (SAR) optimization.

### SAR optimization of CA-01 and CB-01 identifies potent CD28-B7 PPI inhibitors

To improve the potency of the primary hits CA-01 and CB-01, we performed a focused structure-activity relationship (SAR) study by selecting 32 structural analogs for each scaffold, yielding a total of 64 derivatives whose structures are summarized in Table S1 (CA-01 series) and Table S2 (CB-01 series). The 64 derivatives were initially screened at 10 μM against both CD28-CD80 and CD28-CD86 interactions using the NanoBiT assay (Supplementary Fig. S3A-D). This SAR screen identified 5 CA-01-derived analogs and 11 CB-01-derived analogs that conferred significantly enhanced inhibitory activity over the parental compounds. Subsequent dose-response analysis of the most promising derivatives yielded two optimized lead compounds, CA-23 and CB-32, whose chemical structures are shown in Supplementary Fig. S4A, C. CA-23 inhibited CD28-CD80 and CD28-CD86 interactions with IC_50_ values of 35.3 ± 4.3 nM and 23.9 ± 3.1 nM, respectively (Supplementary Fig. S4B), while CB-32 inhibited the same interactions with IC_50_ values of 13.9 ± 3.4 nM and 5.2 ± 2.1 nM, respectively (Supplementary Fig. S4D). These values represent more than a 10-fold improvement in potency over the parental CA-01 and CB-01 scaffolds. CA-23 and CB-32 were therefore selected as the lead compounds for subsequent studies.

### CA-23 binds CD28 selectively over CTLA-4 with a favorable cytotoxicity profile vs. CB-32

A major limitation of the clinically approved CD28-B7 blocker Abatacept is its lack of selectivity, as it engages both CD28 and the inhibitory receptor CTLA-4, thereby compromising regulatory T cell function. To determine whether CA-23 and CB-32 overcome this limitation, we quantitatively assessed their target engagement and selectivity by Microscale Thermophoresis (MST) against recombinant CD28, CD80, CD86, and CTLA-4 proteins. Both compounds bound directly to CD28 with high affinity, with K_D_ values of 0.732 ± 0.016 μM for CA-23 (Fig. 2A) and 0.606 ± 0.01 μM for CB-32 (Fig. 2C). In contrast, Abatacept bound CD80 and CD86 in a dose-dependent manner, with K_D_ values of 0.460 ± 0.124 μM and 0.401 ± 0.074 μM, respectively (Supplementary Fig. S5A, B), consistent with its known mechanism of action as a CTLA-4-Ig fusion that engages B7 ligands rather than CD28 itself. In contrast to this B7-binding profile of Abatacept, neither CA-23 nor CB-32 showed detectable binding to CD80, CD86, or CTLA-4 even at concentrations up to 100 μM (Fig. 2B, D), demonstrating that the two compounds achieve CD28 -specific engagement and discriminate against CTLA-4, a key selectivity gap of Abatacept. Independent surface plasmon resonance (SPR) studies confirmed concentration-dependent CA-23 binding to human and mouse CD28 (K_D_ = 0.35 and 0.52 μM, respectively), whereas CA-23 did not disrupt CTLA-4-CD80 binding in a functional competition assay (Supplementary Fig. S6A,B). No measurable binding of CA-23 to immobilized human CTLA-4 was detected by SPR (Supplementary Fig. S7).

**Fig. 2.**
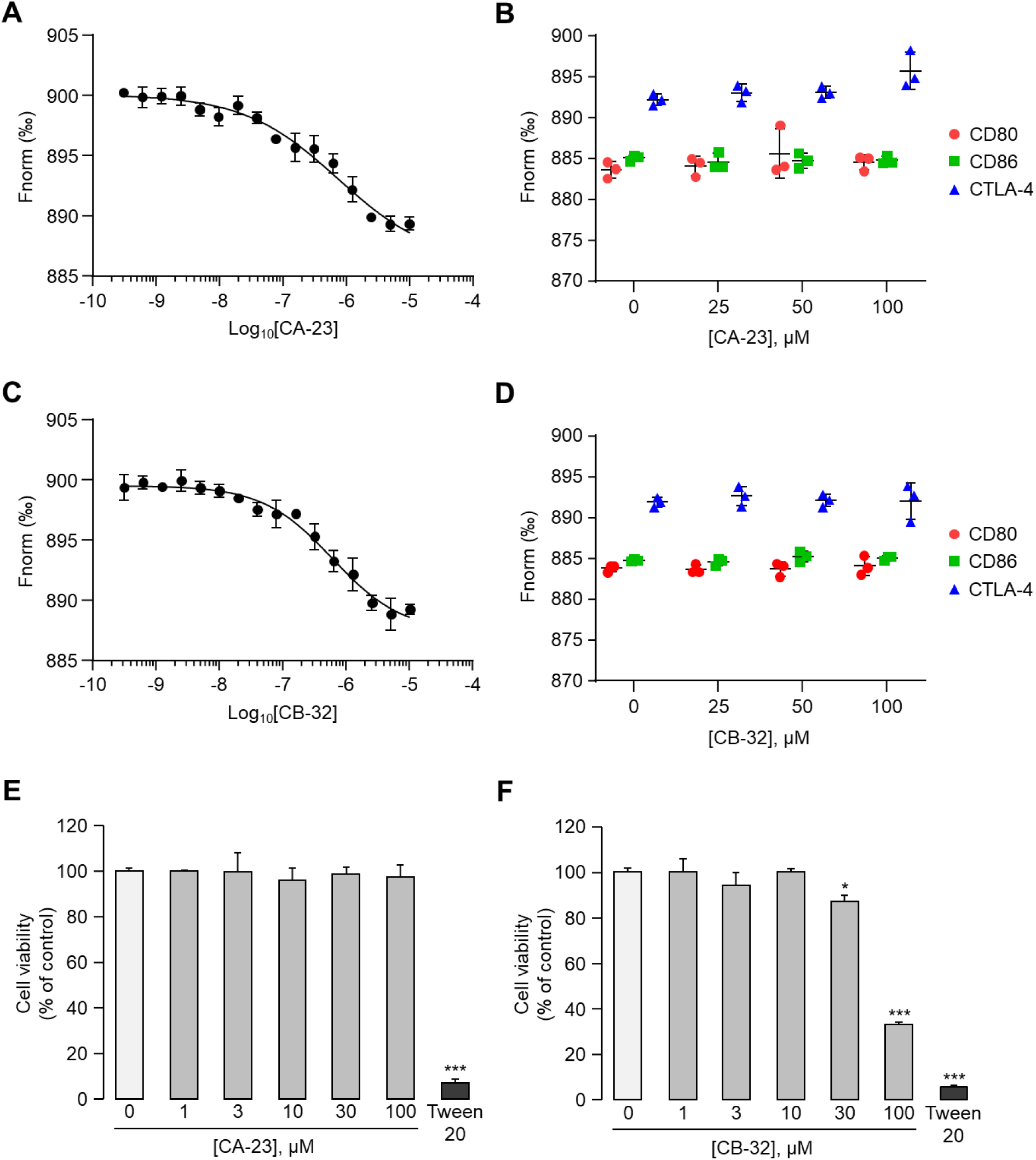
Direct binding of CA-23 and CB-32 to CD28 demonstrates selectivity over CTLA-4 and confirms cell viability. **(A,C)** Microscale Thermophoresis (MST) dose-response curve of CA-23 or CB-32 binding to recombinant CD28 protein. Data are presented as mean ± SD (n=3). **(B,D)** Selectivity profiling of CA-23 **(B)** or CB-32 **(D)** by MST against CD80 (red), CD86 (green), and CTLA-4 (blue) across compound concentrations (0-100 μM). Data are presented as mean ± SD (n=3). **(E, F)** Cell viability of Jurkat cells treated with CA-23 **(E)** or CB-32 **(F)** at the indicated concentrations (0-100 μM) for 24 h. Tween 20 (0.1%) served as a positive control for cytotoxicity. Data are presented as mean ± SD (n=5). *p < 0.05, ***p < 0.001 vs vehicle (one-way ANOVA with Dunnett’s post-hoc test).

To further evaluate the safety profile of the lead compounds, we assessed their cytotoxicity in the T cell-derived Jurkat cell line using the MTS assay. CA-23 exhibited no measurable cytotoxicity up to 100 μM (Fig. 2E), whereas CB-32 caused a modest reduction in viability beginning at 30 μM and pronounced cytotoxicity at 100 μM (Fig. 2F). Although these cytotoxic concentrations are far above the nanomolar IC_50_ values of CB-32 and therefore do not preclude its further development, the combination of potent inhibitory activity, CD28 selectivity, and a clean cytotoxicity profile led us to select CA-23 as the final lead compound for subsequent in vitro and in vivo characterization. CA-23 used in this study was resynthesized in-house with full chemical characterization (^1^H NMR, ^13^C NMR, and LC-MS) provided in Supplementary Information (Supplementary Figure S8).

### CA-23 disrupts CD28-B7 engagement and T cell activation

While the NanoBiT-based assay provides a robust readout of CD28-B7 interaction in a co-expression configuration, it does not faithfully recapitulate the physiologically relevant trans engagement between CD28 on T cells and B7 ligands on antigen-presenting cells (APCs). To overcome this limitation and assess CA-23 in a trans-binding context, we established a series of orthogonal assays. First, we generated HEK293 cells stably expressing CD28 (HEK293-CD28) and examined whether CA-23 could block the engagement of soluble recombinant CD80 or CD86 to surface CD28. HEK293-CD28 cells were pre-treated with CA-23 across a dose range (0-10 μM) for 3 h, followed by incubation with recombinant CD80 (5 μg/ml) or CD86 (10 μg/ml) for 1 h, and cell-bound B7 ligands were quantified by immunoblotting. CA-23 dose-dependently reduced surface binding of both B7 ligands, with detectable inhibition observed at concentrations as low as 100 nM for CD80 (Fig. 3A) and 10 nM for CD86 (Fig. 3B). To more quantitatively characterize the inhibition of the trans CD28-B7 interaction, we performed an enzyme-linked immunosorbent assay (ELISA) using purified recombinant CD28 and biotinylated B7 proteins. In this biochemical assay, residual CD28-B7 binding in the presence of compound was detected via streptavidin-HRP-mediated chemiluminescence, providing a direct readout of trans-like CD28-B7 engagement free from cellular context. CA-23 inhibited the CD28-B7 interaction with an IC_50_ of 180.8 ± 12.8 nM (Fig. 3C), confirming that the disruption is mediated through direct interference with the receptor-ligand interface rather than through cellular co-factors. Finally, we evaluated whether this disruption translates into suppression of CD28-mediated downstream T cell signaling in a fully cellular trans configuration. We co-cultured Jurkat T cells with Raji B cells which serve as B7-expressing antigen-presenting cells and measured CD28-driven activation using a luciferase reporter readout. CA-23 dose-dependently suppressed CD28-mediated T cell activation with an IC_50_ of 424.4 ± 99.3 nM (Fig. 3D). Together, these results demonstrate that CA-23 retains sub-micromolar potency in a trans cellular context that faithfully mirrors the physiological T cell-APC synapse, beyond the co-expression configuration captured by the NanoBiT assay.

**Fig. 3.**
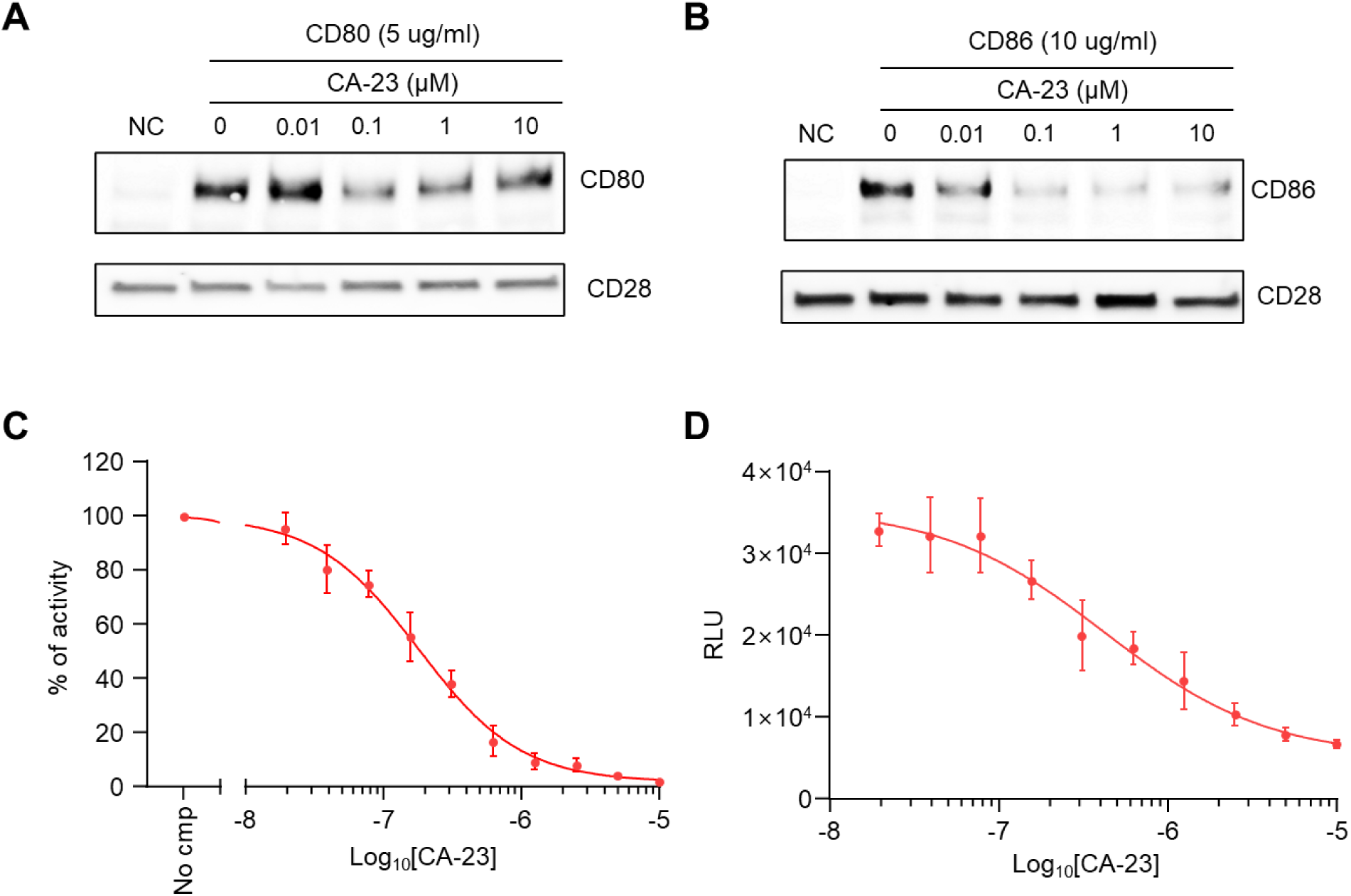
CA-23 inhibits B7 binding to cell-surface CD28 and blocks CD28-mediated T cell activation. **(A, B)** Cell-based binding assay of B7 ligands to surface CD28. HEK293-CD28 cells were pretreated with CA-23 at the indicated concentrations (0-10 μM) for 3 h, followed by incubation with recombinant CD80 protein (5 μg/ml, A) or recombinant CD86 protein (10 μg/ml, B) for 1 h. Cell-bound CD80 or CD86 was detected by immunoblotting (top). CD28 served as a loading control (bottom). **(C)** Inhibition of biochemical CD28-B7 interaction by CA-23 using the CD28:B7 Inhibitor Screening ELISA Kit (BPS Bioscience). Activity is shown as % relative to vehicle (No cmp) control. Data are presented as mean ± SD (n=6). **(D)** Inhibition of CD28-mediated T cell activation by CA-23 in the CD28 Bioassay (Promega). Jurkat-NFAT-luciferase reporter cells were co-cultured with CD80/CD86-expressing artificial APC cells in the presence of CA-23 at the indicated concentrations. NFAT-driven luminescence (RLU) was measured. Data are presented as mean ± SD (n=6).

### CA-23 blocks CD28-driven T cell activation in human blood without independent agonism

Having established potent, CD28-selective inhibition of trans CD28-B7 engagement in cellular and biochemical assays, we next asked whether CA-23 retains this activity in primary human blood and, critically, whether it acts as a pure antagonist without independent agonist activity, a key safety consideration for any CD28-directed agent given the historical precedent of catastrophic cytokine release with CD28 superagonists. In healthy donor PBMCs and whole blood (n=12 donors), CA-23 alone produced no increase in IFN-γ, IL-2, TNF-α, or CD69 expression relative to unstimulated cells, indicating an absence of independent T-cell activation (Fig. 4A). In contrast, CA-23 dose-dependently suppressed TCR/CD28-driven cytokine release back toward TCR-alone levels in both matrices (IFN-γ suppressed to 67%, 39%, and 19% of TCR/CD28-alone levels at 0.1, 0.3, and 1 μM, respectively; Fig. 4B), confirming that CA-23 acts specifically on CD28 co-stimulation rather than TCR signaling itself. We further benchmarked CA-23 potency against FR104, a clinically investigated monovalent anti-CD28 Fab’ antibody with a similar selectivity profile, in primary human CD4^+^ T cells co-cultured with CD80/CD86-expressing artificial antigen-presenting cells. CA-23 inhibited CD28-driven activation with an IC_50_ of 0.28 μM, closely matching the degree of suppression achieved by FR104 at a benchmark 1 μM concentration (Fig. 4C). These data demonstrate that CA-23 functions as a pure CD28 antagonist in human blood and primary human T cells, with potency comparable to a clinical-stage biologic comparator.

**Fig. 4.**
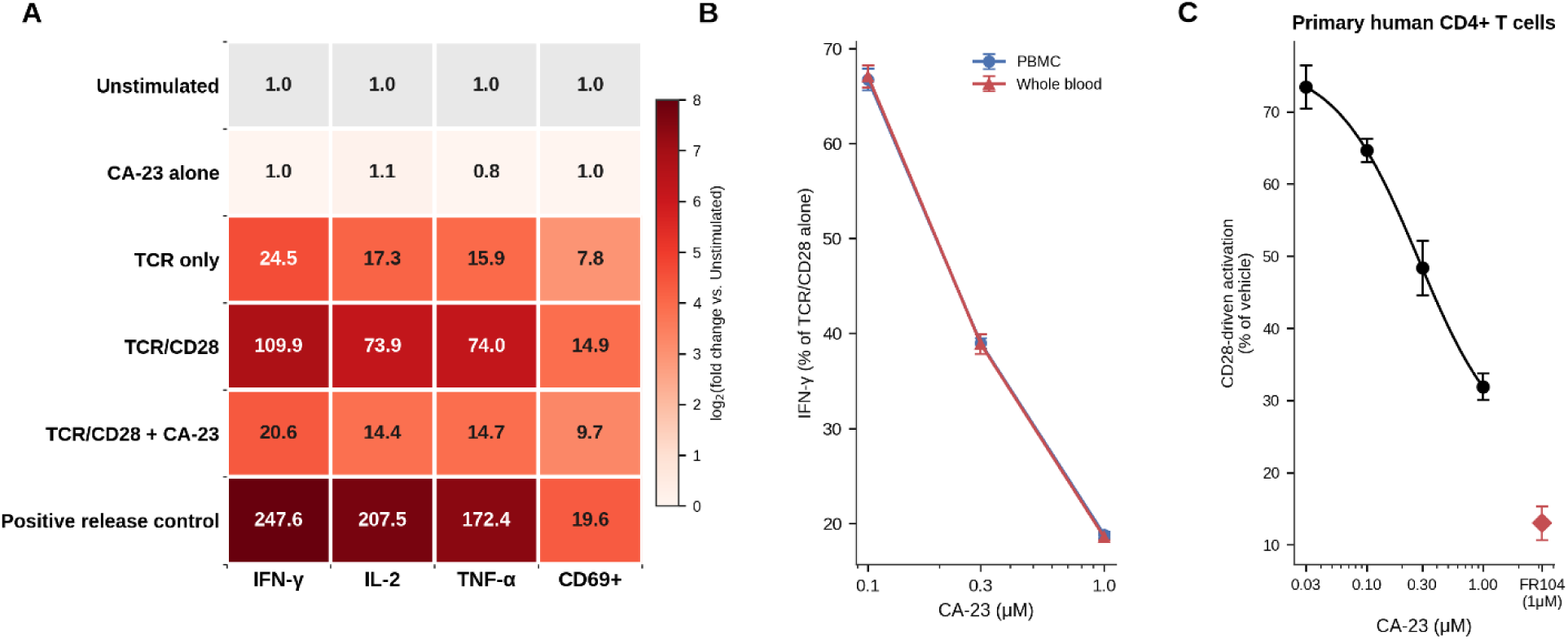
CA-23 blocks CD28-driven T cell activation in human blood without independent agonism. **(A)** Heatmap of cytokine and CD69 responses (fold change vs. unstimulated) in human PBMCs across conditions (n=12 donors). **(B)** Dose-dependent suppression of TCR/CD28-driven IFN-γ by CA-23 in PBMC and whole blood (mean ± SEM). **(C)** Dose-response of CA-23 on CD28-driven activation in primary human CD4^+^ T cells co-cultured with CD80/CD86-expressing artificial antigen-presenting cells; red diamond indicates FR104 benchmark at 1 μM (mean ± SEM).

### In vitro ADME and pharmacokinetics of CA-23 support in vivo evaluation

Before evaluating CA-23 in the colitis animal model, we characterized its in vitro drug-like properties and determined whether the compound achieved exposure in the intestinal and gut-draining lymphoid compartments relevant to disease pathogenesis. CA-23 displayed moderate lipophilicity at physiological pH (LogD_7.4_, 3.1) while retaining favorable kinetic solubility in both aqueous and biorelevant media. Solubility was 85 μM in 1% DMSO/PBS and 152 μM in fasted-state simulated intestinal fluid. CA-23 also showed measurable passive permeability in the PAMPA assay (12.6 × 10^-6^ cm/s), consistent with the potential for passive membrane partitioning.

CA-23 was stable in simulated gastric and intestinal fluids, with 97.2% and 98.1% of compound remaining after 2 hours, respectively. The compound showed moderate metabolic stability in pooled mouse and human liver microsomes, with half-lives of 68.0 and 84.0 minutes, respectively, and remained stable in mouse plasma, human plasma, and PBS throughout the 100-minute assay window. Human plasma protein binding was 94.0%. In early safety studies, CA-23 did not reduce viability of WI-38 or HS-27 cells at concentrations up to 100 μM, showed an hERG IC_50_ of 98.3 μM, and produced less than 15% inhibition across the CYP450, GPCR, and transporter panel at 10 μM (Table S3 and Supplementary Fig. S9). Collectively, these findings supported progression of CA-23 to in vivo pharmacokinetic and efficacy studies.

We next characterized the exposure of CA-23 after a single 10 mg/kg IP dose using the same formulation subsequently used for efficacy studies. CA-23 was rapidly absorbed, reaching a plasma C_max_ of 3.8 μM at 0.5 hours after administration. Plasma exposure declined with a terminal half-life of 3.3 hours and an AUC of 14.2 μM·h.

Importantly, CA-23 distributed to both the colon and mesenteric lymph nodes, the intestinal and gut-draining lymphoid compartments implicated in T-cell transfer colitis. Peak concentrations reached 4.6 nmol/g in colon and 2.1 nmol/g in mesenteric lymph nodes, with colon exposure exceeding plasma exposure on an AUC basis. CA-23 remained detectable in both tissues at 6 hours after administration. These exposure data provided the pharmacokinetic rationale for evaluating daily IP CA-23 administration at 1 and 10 mg/kg in the subsequent colitis efficacy study.

### CA-23 limits T cell transfer-induced colitis and suppresses pathogenic Th1/Th17 responses in vivo

Having established that CA-23 disrupts CD28-B7 engagement and suppresses T cell activation at sub-micromolar potency across multiple orthogonal cellular and biochemical assays, we next examined whether these effects translate into preventive efficacy in a clinically relevant in vivo model of T cell-driven inflammation. We employed the well-established CD4^+^CD45RB^hi^ T cell adoptive transfer model of chronic colitis, in which naïve CD4 ^+^CD45RB^hi^ T cells isolated from the spleens of wild-type donor mice are transferred into Rag1 deficient recipients, where the absence of regulatory T cells permits unrestrained expansion of the transferred T cells and drives progressive intestinal inflammation that recapitulates many features of human IBD.

Mice were treated daily with vehicle, CA-23 (1 mg/kg), or CA-23 (10 mg/kg), and body weight was monitored every three days as a global indicator of disease progression. CD4^+^CD45RB^hi^ T cell-transferred mice treated with vehicle exhibited significant weight loss beginning at day 24 post-transfer, consistent with the established disease kinetics of this model (Fig. 5A). In contrast, CA-23 treatment dose-dependently attenuated weight loss: mice receiving 1 mg/kg only showed mild divergence from negative control animals near the experimental endpoint (day 42), whereas mice injected CA-23 10 mg/kg maintained body weight indistinguishable from negative controls throughout the entire observation period.

**Fig. 5.**
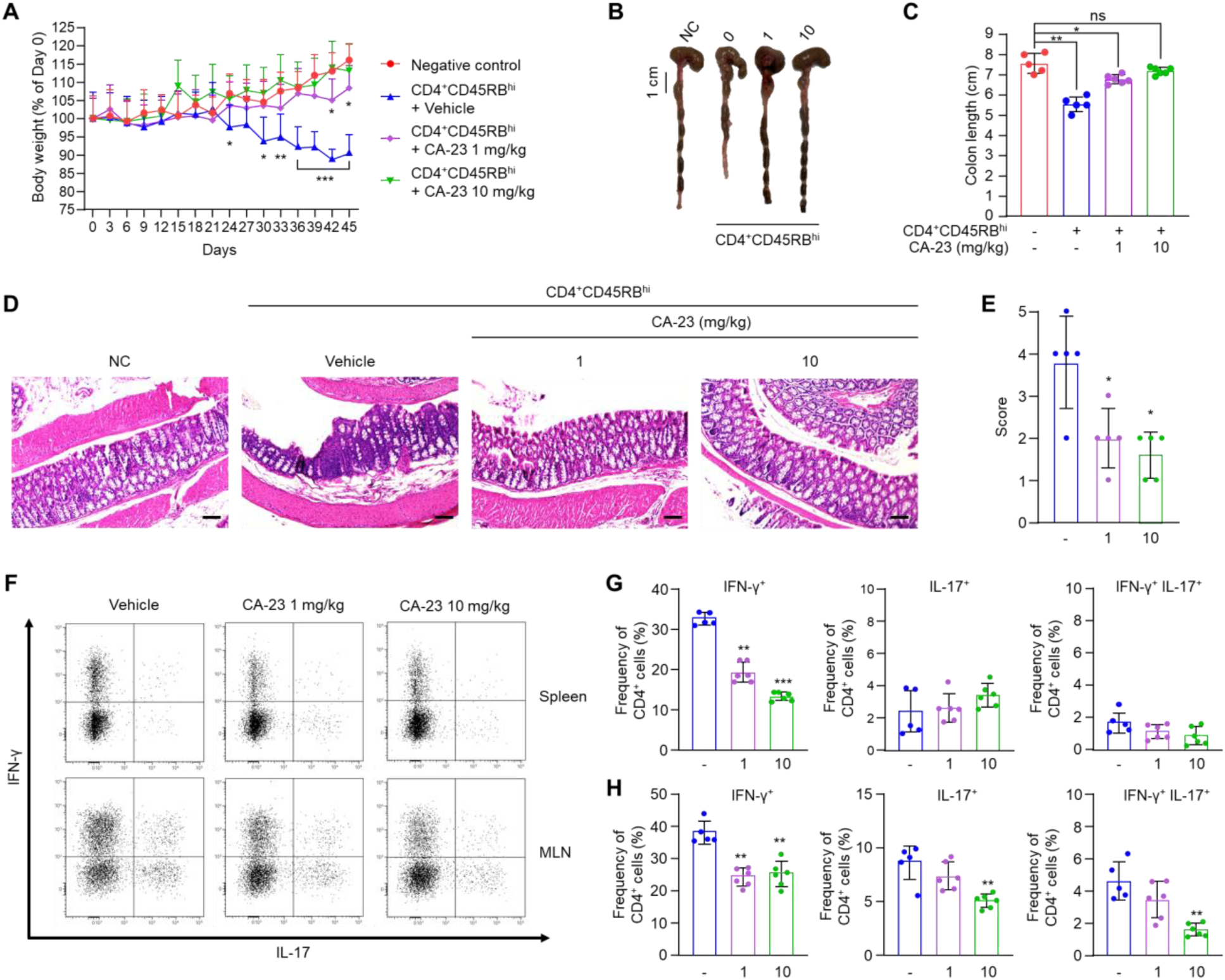
CA-23 ameliorates T cell transfer-induced colitis by suppressing pathogenic Th1 and Th17 responses in vivo. (A) Body weight changes of Rag1^-/-^ mice over 45 days following adoptive transfer of CD4^+^CD45RB^hi^ T cells. Mice were treated with vehicle, CA-23 (1 mg/kg), or CA-23 (10 mg/kg). Negative control mice received no T cell transfer. Body weight is shown as % of Day 0. Data are presented as mean ± SD (n=5-7 per group). *p < 0.05, **p < 0.01, ***p < 0.001 vs Negative control at the same time point (two-way ANOVA with Tukey’s post-hoc test). (B) Representative images of colons at endpoint (Day 45). Scale bar, 1 cm. NC, negative control; 0, vehicle; 1, CA-23 1 mg/kg; 10, CA-23 10 mg/kg. (C) Quantification of colon length. Data are presented as mean ± SD (n=5-6 per group). *p < 0.05, **p < 0.01; ns, not significant (one-way ANOVA with Tukey’s post-hoc test). (D) Representative H&E-stained sections of colonic tissue from each group. Scale bars (bottom right of each image), 100 μm. (E) Histological scoring of colonic inflammation performed in a blinded manner. Data are presented as mean ± SD (n=5 per group). *p < 0.05 vs vehicle (one-way ANOVA with Tukey’s post-hoc test). (F) Representative flow cytometry plots of IFN-γ and IL-17A expression in CD4^+^ T cells from spleen (top) and MLN (bottom). Cells were stimulated ex vivo with PMA/ionomycin in the presence of brefeldin A for 5 h prior to intracellular cytokine staining. (G) Frequencies of IFN-γ^+^, IL-17^+^, and IFN-γ^+^IL-17^+^ CD4^+^ T cells in the spleen of CD4^+^CD45RB^hi^ T cell-transferred mice treated with vehicle (-), 1 (1 mg/kg CA-23), or 10 (10 mg/kg CA-23). Data are presented as mean ± SD (n=5-6 per group). **p < 0.01, ***p < 0.001 vs vehicle (one-way ANOVA with Tukey’s post-hoc test). (H) Frequencies of IFN-γ^+^, IL-17^+^, and IFN-γ^+^IL-17^+^ CD4^+^ T cells in the MLN, displayed as in (G). Data are presented as mean ± SD (n=5-6 per group). **p < 0.01 vs vehicle (one-way ANOVA with Tukey’s post-hoc test).

To complement the systemic readout, we assessed colonic pathology at endpoint. Shortening of the colon is a well-established hallmark of chronic intestinal inflammation. Compared with negative control animals, vehicle-treated colitic mice exhibited markedly shortened colons, which were progressively restored by CA-23 in a dose-dependent manner, with 10 mg/kg fully normalizing colon length to negative control levels (Fig. 5B, C). Histopathological examination of H&E-stained colonic sections revealed crypt distortion, epithelial erosion, goblet cell loss, and transmural infiltration of inflammatory cells in vehicle-treated colitic mice, recapitulating features typically reported in CD4^+^CD45RB^hi^-induced colitis (Fig. 5D). CA-23 treatment dose-dependently preserved crypt architecture, restored goblet cell content, and reduced inflammatory cell infiltration, leading to significantly lower histopathological scores compared with vehicle-treated mice (Fig. 5E).

To dissect the immunological mechanism underlying this protective effect, we analyzed CD4 ^+^ T cell cytokine profiles by intracellular flow cytometry in the spleen and mesenteric lymph nodes (MLN), the secondary lymphoid organ that drains the colonic mucosa (Fig. 5F). In the spleen, IL-17^+^ CD4^+^ T cells were sparse across all groups, whereas IFN-γ^+^ CD4^+^ T cells were markedly enriched in vehicle-treated colitic mice (∼32% of CD4^+^ cells), consistent with the historical characterization of CD45RB^hi^-driven colitis as a Th1-dominated systemic response. CA-23 dose-dependently and significantly reduced this Th1 expansion to ∼19% at 1 mg/kg and ∼13% at 10 mg/kg (Fig. 5G).

In the MLN, in addition to abundant IFN-γ^+^ cells, IL-17^+^ CD4^+^ T cells were observed at higher frequencies than in the spleen (∼10%), along with a detectable population of IFN-γ^+^IL-17^+^ double-positive cells. Similar enrichment of Th17 and double-positive populations in gut-draining lymphoid tissues has been reported in the CD45RB^hi^ colitis model, and double-positive CD4^+^ T cells have likewise been described in intestinal tissues of IBD patients. The pattern observed in our MLN is consistent with these previous reports, indicating that the model faithfully recapitulates the gut-local Th1/Th17 immune environment described in human IBD. Against this pathogenic immune backdrop, CA-23 significantly reduced IFN-γ^+^ CD4^+^ T cells in the MLN at both 1 mg/kg and 10 mg/kg, and significantly reduced IL-17^+^ and IFN-γ^+^IL-17^+^ double-positive populations at 10 mg/kg (Fig. 5H).

Collectively, these data demonstrate that CA-23 confers significant in vivo preventive efficacy in a T cell-driven colitis model. Importantly, CA-23 suppressed pathogenic Th1 responses across both systemic and mucosal-draining compartments and further suppressed Th17 and IFN-γ^+^IL-17^+^ double-positive populations at the MLN in a dose-dependent manner, thereby broadly targeting the key pathogenic immune environment that develops at the disease-proximal lymphoid tissue.

### CA-23 suppresses pathogenic cytokine and Th1/Th17 responses in PBMCs from patients with active IBD

To extend these findings to human disease-relevant material, we obtained PBMCs from 30 patients with active IBD (15 ulcerative colitis (UC), 15 Crohn’s disease) and evaluated CA-23 head-to-head against FR104 and Abatacept at a matched top concentration (1 μM). CA-23 dose-dependently suppressed IFN-γ, IL-2, TNF-α, IL-17A, and GM-CSF in both disease groups (Fig. 6A, B), with a magnitude of suppression comparable to Abatacept on four of the five cytokines (e.g., IFN-γ suppressed to 47% and 45% of vehicle by CA-23 in UC and Crohn’s PBMCs, respectively, versus 48% and 39% by Abatacept), while FR104 produced the greatest suppression of the three agents. IL-17A was the exception: CA-23 suppressed IL-17A less than Abatacept in both disease groups (61% vs. 43-44% of vehicle; q=0.03 UC, q=0.0003 Crohn’s, paired t-test with FDR correction), the only cytokine where the two agents diverged significantly. Flow cytometric analysis of CD4^+^ T-cell subsets corroborated this pattern: CA-23 suppressed CD69^+^, Ki67^+^, IFN-γ^+^, IL-17A^+^, and IFN-γ^+^IL-17A^+^ double-producing CD4^+^ T-cell frequencies significantly more than Abatacept in both UC and Crohn’s PBMCs (all q≤0.002, paired t-test with FDR correction; Fig. 6C), with FR104 again producing the greatest suppression of the three agents. Notably, FOXP3^+^ Treg frequency was unchanged across all four conditions (vehicle, CA-23, FR104, Abatacept) in this short-term assay, indicating that none of the three agents altered Treg abundance over this timeframe, a finding that motivated the direct functional assessment of Treg activity described below.

**Fig. 6.**
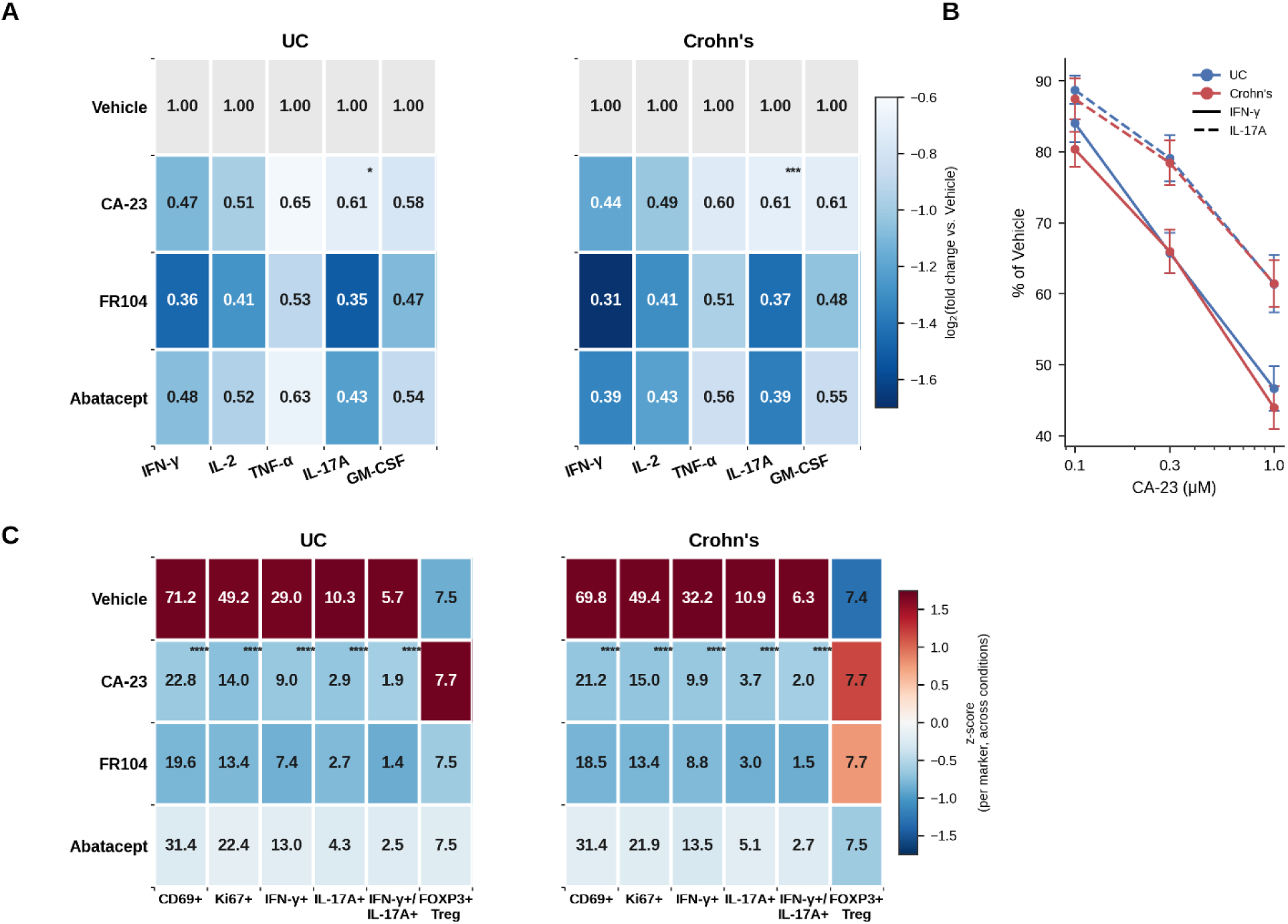
CA-23 suppresses pathogenic cytokine and Th1/Th17 responses in PBMCs from IBD patients. **(A)** Heatmap of cytokine responses (fold change vs. donor-matched Vehicle) at matched top concentration (1 μM/equivalent) in UC and Crohn’s patient PBMCs (n=15 donors/group); Vehicle shown as fixed reference (=1.00). *q<0.05, ***q<0.001, CA-23 vs. Abatacept (paired t-test, Benjamini-Hochberg FDR-corrected within disease group). **(B)** Dose-dependent suppression of IFN-γ and IL-17A by CA-23 (mean ± SEM). **(C)** Heatmap of CD4^+^ T-cell subset frequencies by flow cytometry (n=15 donors/group), including FOXP3^+^ Treg frequency; color denotes z-score within each marker across conditions. ****q<0.0001, CA-23 vs. Vehicle (effector markers); CA-23 vs. Abatacept also significant on all five effector markers (q≤0.002) and non-significant on FOXP3^+^ Treg (paired t-test, FDR-corrected).

### CA-23 preserves regulatory T cell function and epithelial barrier integrity in vitro

A central hypothesis of this work is that CD28-selective antagonism, by sparing CTLA-4, should preserve Treg function where Abatacept does not. We tested this directly using an autologous Treg suppression assay in 12 IBD patient donors (6 UC, 6 Crohn’s). Treg suppressive capacity was unchanged by CA-23 or FR104 relative to vehicle (UC: 52.0% vehicle vs. 53.4% CA-23 vs. 51.6% FR104; Crohn’s: 54.8% vs. 54.9% vs. 56.6%), whereas Abatacept nearly halved Treg suppressive capacity in both disease groups (32.2% UC, 36.1% Crohn’s; p<0.001 vs. vehicle and vs. CA-23 in both groups; Fig. 7A), consistent with preservation of Treg suppressive function during CD28-selective antagonism at the functional level. In the same assay, CA-23 dose-dependently suppressed responder T-cell proliferation (Fig. 7B), confirming on-mechanism efficacy in parallel with preserved Treg function. Using a human intestinal epithelial–PBMC co-culture system, CA-23 improved trans-epithelial electrical resistance (TEER) and reduced FITC-dextran permeability relative to vehicle, comparable to FR104 and Abatacept (Fig. 7C, D), indicating protection of epithelial barrier integrity, a property of direct mechanistic relevance to IBD. Finally, in an autologous antigen-presenting cell–T-cell co-culture system, CA-23 suppressed CD4⁺ T-cell activation across its dose range without altering CD80 or CD86 surface expression on autologous monocytes (Fig. 7E), showing that CA-23 did not alter APC B7-ligand expression.

**Fig. 7.**
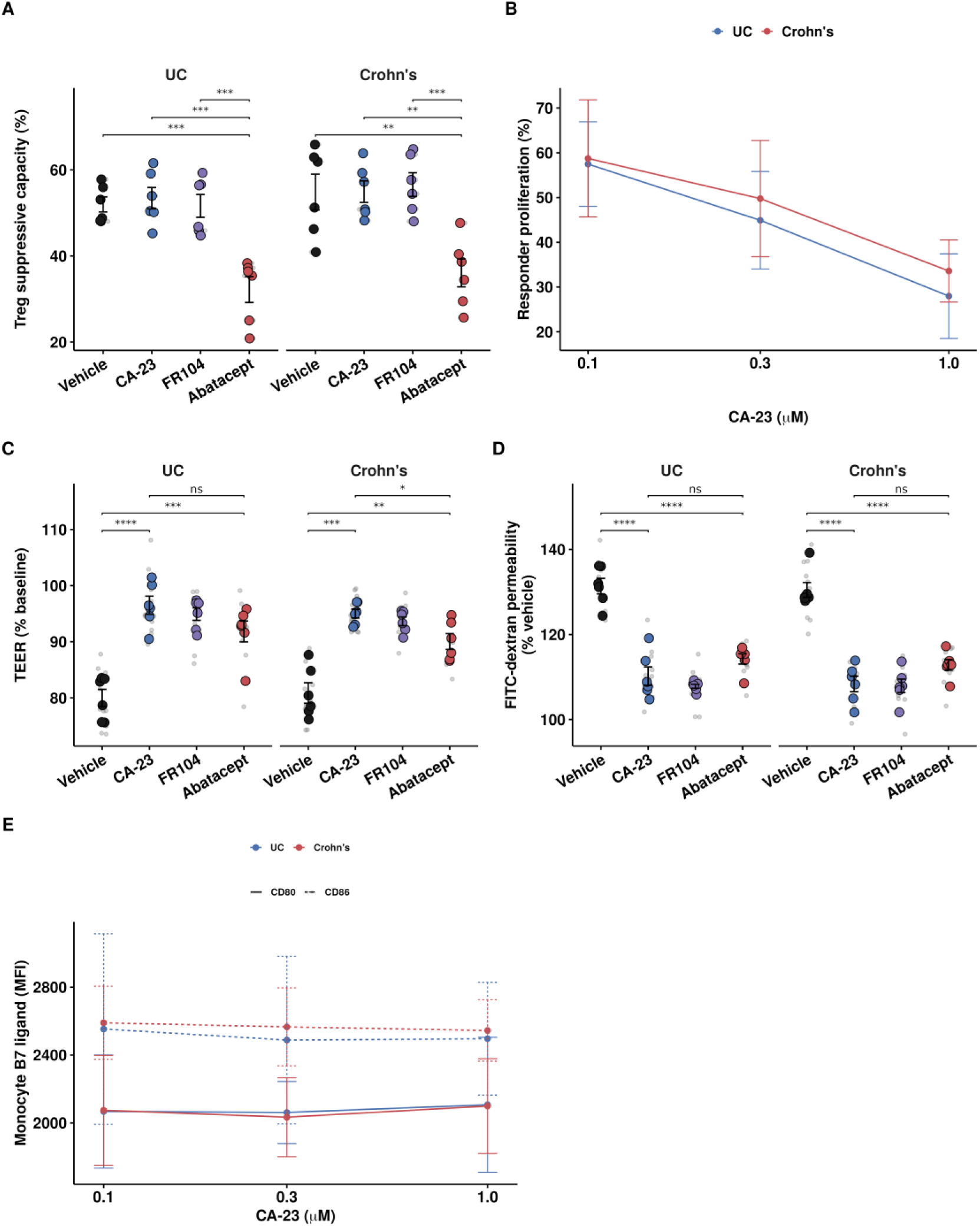
CA-23 preserves regulatory T cell function and epithelial barrier integrity in vitro. **(A)** Treg suppressive capacity in an autologous Treg suppression assay (n=6 donors/disease group); each large point is a donor mean, small points are technical replicates. **(B)** CA-23 dose-response of responder T-cell proliferation. **(C, D)** Epithelial barrier TEER **(C)** and FITC-dextran permeability **(D)** in human intestinal epithelial–PBMC co-cultures. **(E)** CD80 and CD86 surface expression (MFI) on autologous monocytes across CA-23 doses. Data in A, C, D are shown as donor-level means ± SEM with statistics on donor means (t-test); data in B, E are mean ± SD.

## DISCUSSION

In this study, we developed a sensitive HiBiT-LgBiT NanoBiT split-luciferase platform to enable high-throughput identification of direct small molecule antagonists of the CD28 -B7 protein-protein interaction, a historically intractable immunotherapeutic target. Screening of approximately 5,000 structurally diverse compounds, followed by focused structure-activity relationship optimization of two distinct hit scaffolds, yielded CA-23, a CD28-selective small molecule antagonist with nanomolar potency in cellular CD28-B7 interaction assays. CA-23 binds CD28 directly with sub-micromolar affinity, exhibits no detectable engagement of CD80, CD86, or CTLA-4 even at 100 μM, and inhibits trans CD28-B7 binding and downstream T cell activation across multiple orthogonal cellular and biochemical assays. In a CD4 ^+^CD45RB^hi^ T cell adoptive transfer model of chronic colitis, CA-23 dose-dependently ameliorated weight loss, restored colon length, reduced histopathological inflammation, and suppressed pathogenic Th1, Th17, and IFN - γ^+^IL-17A^+^ double-producing CD4^+^ T cells at the disease-proximal mesenteric lymph nodes. Together, these findings establish a CD28-selective small molecule antagonist with experimentally validated selectivity over CTLA-4 and demonstrated in vivo efficacy in a T cell-driven model of IBD.

A key methodological advance of this work was the application of the HiBiT-LgBiT NanoBiT system as a cell-based HTS assay for the CD28-B7 interaction. The high-affinity HiBiT variant (K_D_ ≈ 700 pM, versus ≈190 μM for conventional SmBiT) substantially improves signal brightness and dynamic range, enabling robust detection of weak and transient receptor-ligand contacts characteristic of many co-signaling PPIs (*27*, *28*). The high Z’ factors obtained for both the CD28-CD80 (Z’ = 0.756) and CD28-CD86 (Z’ = 0.621) assays exceeded the conventional HTS threshold, confirming the platform’s suitability for primary compound screening (*30*). Performing the screen in a cellular context ensured hits disrupted CD28-B7 engagement in its native membrane environment, while parallel counter-screening against an unrelated NOS1-NOS1AP NanoBiT pair excluded promiscuous and assay-interfering compounds early (*29*). Although demonstrated here only for CD28-B7, this HiBiT-LgBiT configuration may extend to other weak or transient co-signaling PPIs, pending target-by-target validation.

Beyond the in vivo colitis model, a central goal of this work was to establish the translational relevance of CA-23 and define its pharmacokinetic and safety properties. CA-23 acted as a pure CD28 antagonist in human PBMCs and whole blood, with no independent T-cell agonism and dose-dependent suppression of CD28-driven cytokine release, and achieved tissue exposure in colon and MLN supporting local pharmacodynamic activity. In PBMCs from patients with active UC or Crohn’s disease, CA-23 matched or exceeded Abatacept on pathogenic cytokine and Th1/Th17 suppression, significantly outperforming it on IL-17A and on all five flow cytometric effector markers, while directly demonstrating, through a functional Treg suppression assay, that CA-23 preserves regulatory T-cell function where Abatacept does not. This functional dissociation is, to our knowledge, the first direct experimental confirmation in human disease-relevant cells that a CD28-selective small molecule can match or exceed a B7-directed biologic on efficacy-relevant readouts while avoiding its principal mechanistic liability.

The most distinctive feature of CA-23 is its direct and selective engagement of CD28 without measurable binding to CD80, CD86, or CTLA-4. This selectivity profile addresses the principal mechanistic liability of Abatacept, the only clinically approved co-stimulation blocker. Because Abatacept is a CTLA-4-Ig fusion that binds CD80 and CD86 on antigen-presenting cells, it indiscriminately occupies B7 ligands and disrupts both the stimulatory CD28 -B7 and the inhibitory CTLA-4-B7 axes (*17*, *18*, *23*, *24*). By binding CD28 directly and sparing CTLA-4, CA-23 selectively disrupts the stimulatory arm of the CD28-B7 axis while leaving the CTLA-4-B7 interaction intact, a mechanism conceptually distinct from B7 sequestration and aligned with the design principle of selective anti-CD28 biologics such as FR104 (*35*). We have previously contributed to validating the chemical tractability of CD28 through the discovery of small molecule CD28 binders identified by affinity selection-mass spectrometry, structure-based virtual screening, and biophysical fragment approaches (*36*, *37*). The present work advances this effort by introducing CA-23, a CD28 antagonist that, unlike previously reported small molecule binders, has been explicitly validated for CTLA-4 selectivity by direct biophysical measurement and shown to be efficacious in an in vivo model of T cell-driven disease. This selectivity was independently corroborated by SPR, which showed direct CA-23 binding to human and mouse CD28 and no measurable binding to human CTLA-4, and by a functional CTLA-4-CD80 competition assay. The similar affinity of CA-23 for human and mouse CD28 supports the translatability of the in vivo efficacy findings described below. Functionally, this CTLA-4-sparing selectivity translated directly into preserved Treg suppressive capacity in patient-derived cells, in contrast to the marked impairment of Treg function observed with Abatacept in the same assay. Co-crystallographic or cryo-EM analysis of the CA-23-CD28 complex will still be required to define the binding site at atomic resolution and the precise structural basis of this selectivity.

The in vivo activity of CA-23 showed a compartment-specific pattern. In the spleen, IL-17A-producing Th17 cells were sparse across all groups, while IFN-γ-producing Th1 cells dominated the colitic state and were dose-dependently suppressed by CA-23. In the mesenteric lymph nodes (MLN), the principal site of pathogenic T cell expansion in this model, Th17 cells and IFN -γ^+^IL-17A^+^ double-producing CD4^+^ T cells accumulated to higher frequencies than in the spleen, and CA-23 suppressed these subsets at 10 mg/kg. This is consistent with the known compartmentalization of effector T cells in chronic intestinal inflammation, where Th17 and double-producing cells preferentially expand at gut-associated lymphoid tissues under IL-23-rich mucosal cytokine milieus, while Th1-skewed responses predominate systemically (*13*, *14*). This double-producing population has been repeatedly identified in inflamed Crohn’s mucosa and chronic colitis models as a highly pathogenic effector subset arising through Th17-to-Th1 plasticity (*12*, *14*); the CD4^+^CD45RB^hi^ transfer model used here, with its Th1-dominant systemic phenotype and mucosa-enriched Th17/double-producing populations, recapitulates key immunological features of Crohn’s disease (*31*). This ability to suppress both the MLN-enriched double-producing population and systemic Th1 expansion is consistent with upstream co-stimulation blockade broadly attenuating the pathogenic effector T cell network, in contrast with the single-cytokine neutralization of currently approved biologics such as anti-TNF-α or anti-IL-12/23 antibodies, raising the possibility that CD28-selective antagonism may benefit patients with primary non-response or secondary loss of response to single-cytokine-targeted biologics (*15*, *16*). Notably, Abatacept itself was evaluated in clinical trials for Crohn’s disease and ulcerative colitis but failed to demonstrate meaningful efficacy (*38*). This contrast between Abatacept’s clinical failure via B7 sequestration and the in vivo efficacy of CA-23 observed here raises the possibility that the two mechanisms may yield differential outcomes in IBD, a hypothesis warranting systematic head-to-head testing in matched preclinical models.

The findings presented here suggest several productive directions for future investigation. First, although CA-23 binds CD28 directly with sub-micromolar affinity by both MST and SPR and shows complete selectivity against CD80/CD86/CTLA-4, its precise binding site has not yet been mapped; co-crystallographic or cryo-EM characterization of the CA-23-CD28 complex will define the structural basis of this selectivity and support further potency and drug-like property optimization. Second, this in vitro comparison with Abatacept has not yet been extended to a matched in vivo IBD model; given that Abatacept itself failed in clinical trials for Crohn’s disease and ulcerative colitis (*38*), a head-to-head comparison in the CD4+CD45RBhi transfer model or an independent colitis model would help establish whether CA-23’s Treg-preserving mechanism translates into superior in vivo efficacy. Finally, given that CD28-mediated co-stimulation plays a central pathogenic role across T cell-driven immune-mediated diseases broadly (*8*), CA-23’s therapeutic potential is unlikely to be confined to the Crohn’s-like IBD captured here. Rheumatoid arthritis, Abatacept’s principal approved indication (*17*, *18*), and transplant rejection, in which the related CTLA-4-Ig fusion Belatacept is clinically established (*39*), represent natural next adjacencies for CA-23 evaluation in disease-specific immune environments such as collagen-induced arthritis (*40*) and experimental autoimmune encephalomyelitis (*41*).

In summary, we report a CD28-selective small molecule antagonist, CA-23, that combines direct, SPR-validated CD28 engagement with no measurable CTLA-4 binding, favorable pharmacokinetics with target-proximal tissue exposure, clean in vitro selectivity, and demonstrated in vivo efficacy in a T cell-driven model of IBD. In PBMCs and functional co-culture assays from patients with active IBD, CA-23 matched or exceeded Abatacept on pathogenic cytokine and Th1/Th17 suppression while preserving regulatory T-cell function and epithelial barrier integrity, validating the mechanistic hypothesis that CD28-selective antagonism can dissociate efficacy from the Treg-impairing liability of B7-directed biologics. Together with the HiBiT-LgBiT NanoBiT platform established here, this work provides a foundation for next-generation, mechanism-differentiated CD28-directed immunomodulatory therapies for IBD and other chronic T cell-driven inflammatory diseases.

## MATERIALS AND METHODS

### Cell culture and transfections

CHO-K1 and Jurkat cells were cultured in RPMI 1640 medium supplemented with 10% fetal bovine serum (FBS), 100 U/ml penicillin, and 100 μg/ml streptomycin at 37 ℃ in a humidified incubator with 5% CO₂. For NanoBiT-based screening of CD28-B7 protein-protein interaction (PPI) inhibitors, CHO-K1 cells were co-transduced with lentiviral vectors (pLenti6P.3) encoding HiBiT-CD28 and either LgBiT-CD80 or LgBiT-CD86. Stable cell lines were selected with 1 μg/ml puromycin and 10 μg/ml blasticidin. CHO-K1 cells stably co-expressing the NOS1-NOS1AP NanoBiT pair, used for counter-screening, were generated as described in our previous study (*29*).

For cell-based binding and immunoblot assays, HEK293 cells stably expressing CD28 (HEK293-CD28) were generated using the same lentiviral system and maintained in DMEM supplemented with 10% FBS and 1% penicillin/streptomycin.

### Compounds and reagents

Compound sources and catalog details are provided in the Supplementary Materials and Methods.

### NanoBiT-based high-throughput screening and dose-response assays

CHO-K1 cells stably expressing HiBiT-CD28 and LgBiT-CD80 or LgBiT-CD86 were seeded in 96-well plates and grown to approximately 80% confluency. Cells were washed twice with PBS and overlaid with 100 μl of HBSS solution (140 mM NaCl, 5 mM KCl, 1 mM MgCl2, 1 mM CaCl2, 10 mM D-glucose, 10 mM HEPES, pH 7.4 adjusted with NaOH). Test compounds were added at the indicated concentrations and incubated for 3 h at 37 ℃. Furimazine (final concentration 10 μM) was then added, and plates were incubated for an additional 10 min at 37 ℃ before luminescence (460 nm) was measured on an Infinite M1000 Pro Microplate Reader (Tecan, Männedorf, Switzerland). Luminescence values were normalized as a percentage of DMSO-treated control wells (100%) and parental CHO-K1 cells lacking the NanoBiT constructs (0%). For primary HTS of the Discovery Diversity Set, compounds were pooled four per well (4 -in-1 format) at a final concentration of 25 μM per compound. Hits were defined as exhibiting greater than 70% reduction in luminescence relative to DMSO control. Counter-screening against the NOS1-NOS1AP NanoBiT pair was performed under identical conditions, as previously described (*29*). Dose-response curves were generated using indicated concentration and IC50 values were calculated by four-parameter logistic regression using GraphPad Prism 10 (GraphPad Software, San Diego, CA, USA).

### Z’ factor determination

Assay robustness was evaluated by calculating the Z’ factor using the following formula: Z’ = 1 − [3 × (σpos + σneg) / |μpos − μneg|], where σ and μ represent the standard deviation and mean of positive (Abatacept treated) and negative (DMSO treated) controls, respectively (*30*). A Z’ factor greater than 0.5 was considered indicative of a high-quality assay suitable for HTS.

### Microscale Thermophoresis (MST) binding assay

Binding affinity was measured using a Monolith X instrument (NanoTemper Technologies, Munich, Germany). Recombinant human CD28, CD80, CD86, and CTLA-4 proteins were labeled with the RED-tris-NTA His-tag labeling kit (NanoTemper Technologies) per manufacturer’s instructions. Labeled proteins (40 nM final) were incubated with serially diluted test compounds in PBS containing 0.05% Tween-20 (pH 7.4) for 10 min at room temperature. MST measurements were performed using standard capillaries with 100% LED power and medium MST power; thermophoresis was monitored for 10 s with a 5-s delay. Data were processed using MO. Affinity Analysis software (NanoTemper Technologies) and K_D_ values were determined by curve fitting in GraphPad Prism 10.

### Cell viability assay

The cytotoxic effects of compounds were evaluated using the CellTiter 96® AQueous One Solution Cell Proliferation Assay (Promega, Madison, WI, USA). Jurkat T cells were seeded in 96-well plates (1 × 10⁵ cells/well) and treated with compounds at the indicated concentrations for 24 h. Tween-20 (0.1%) was included as a positive control for cytotoxicity. MTS reagent was added per manufacturer’s protocol, and absorbance was measured at 490 nm using an Infinite M1000 Pro Microplate Reader (Tecan). Cell viability was expressed as a percentage of vehicle (DMSO) treated control wells.

### Cell based B7 binding assay and immunoblotting

HEK293-CD28 cells were seeded in 6-well plates and incubated overnight. Cells were pretreated with CA-23 at the indicated concentrations for 3 h, followed by incubation with recombinant CD80 (5 μg/ml) or CD86 (10 μg/ml) for 1 h. Cells were then washed three times with ice-cold PBS and lysed for 15 min on ice in RIPA buffer supplemented with a protease inhibitor cocktail. Lysates were cleared by centrifugation (13,000 rpm, 20 min, 4 ℃), and protein concentrations were determined using the Bradford assay (Thermo Fisher Scientific, Waltham, MA, USA). Total protein (40 μg per lane) was separated on 4-20% Tris-glycine precast gels (Bio-Rad, Hercules, CA, USA) and transferred to PVDF membranes (Bio-Rad). Membranes were blocked for 1 h with SuperBlock™ Blocking Buffer (Thermo Fisher) and incubated overnight at 4 ℃ with primary antibodies against CD28 (Cell Signaling Technology, Cat. #38774, 1:1000), CD80 (Cell Signaling Technology, Cat. #15416, 1:1000), or CD86 (Cell Signaling Technology, Cat. #91882, 1:1000). After washing with TBST, membranes were incubated with HRP-conjugated secondary antibodies (1:5000) for 1 h at room temperature, and signals were detected using the SuperSignal™ West Pico PLUS Chemiluminescent Substrate (Thermo Fisher).

### Competitive ELISA Analysis of CD28–CD80 Interaction Blockade

Competitive ELISA experiments were carried out using the CD28 [Biotinylated] Inhibitor Screening Kit (BPS Bioscience, Cat. #72007) per manufacturer’s protocol with minor modifications. Briefly, recombinant human CD28 protein was diluted to 2 µg/mL in PBS and immobilized onto 96-well plates (50 µL/well) by overnight incubation at 4 ℃. Plates were subsequently washed with 1× Immuno Buffer and blocked with Blocking Buffer for 1 h at room temperature.

To evaluate inhibitory activity, serial dilutions of test compounds were pre-incubated with biotinylated CD80 (5 ng/µL) for 1 h at room temperature before being transferred to CD28-coated wells. Wells lacking immobilized CD28 served as ligand-only controls, while wells containing inhibitor buffer without compound were used as negative controls. Following incubation and washing steps, streptavidin-conjugated HRP diluted 1:1000 in Blocking Buffer was added and incubated for 1 h at room temperature. Chemiluminescent substrate was then added, and luminescence signals were recorded immediately using an Infinite M1000 Pro microplate reader (Tecan, Switzerland). Concentration–response curves were generated using GraphPad Prism 10, and IC_50_ values were calculated by four-parameter logistic regression analysis. All experiments were performed with the number of replicates indicated in the corresponding figure legends.

### Functional Inhibition of CD28 Signaling Using a Cell-Based Reporter Assay

Functional blockade of CD28 signalling was evaluated using the CD28 Blockade Bioassay kit (Promega, Cat. #JA6101) per manufacturer’s recommendations with minor modifications. Briefly, Jurkat CD28 Effector Cells were seeded into white 96-well plates at a density of 2 × 10^4^ cells per well and pre-treated for 5 min with serial dilutions of test compounds prepared as a 10-point 1:1 dilution series starting from 200 µM with a final DMSO concentration of 1%. An anti-CD28 control antibody (Promega, Cat. #K1231) was included as a positive inhibition control.

Following compound pre-incubation, aAPC/Raji Cells were added at 2 × 10^4^ cells per well, and the plates were incubated for 5 h at 37 ℃ in a humidified atmosphere containing 5% CO₂. Subsequently, Bio-Glo™ Luciferase Reagent (Promega) was added per manufacturer’s protocol, and luminescence signals were recorded using a GloMax® Discover System (Promega). Dose– response curves were analyzed using GraphPad Prism 10 with a four-parameter logistic regression model to determine IC_50_ values. All experiments were conducted in triplicate, and data are presented as mean ± SD. The assay maintained robust performance in the presence of up to 10% pooled human serum, indicating compatibility with physiologically relevant assay conditions.

### Human PBMC samples

Cryopreserved PBMCs from donors with UC or Crohn’s disease were obtained from a commercial supplier (Sanguine Biosciences) as deidentified research-use material. All donors had documented IBD in the supplier-provided records. Samples were therefore classified by reported diagnosis only. Fifteen UC and fifteen Crohn’s disease donors were used for the core PBMC cytokine and flow-cytometry experiments.

PBMCs were thawed rapidly in a 37°C water bath, transferred dropwise into prewarmed RPMI 1640 supplemented with 10% heat-inactivated FBS, 1% penicillin-streptomycin, and benzonase (50 U/mL), and washed twice. Cells were rested overnight in complete RPMI at 37°C in 5% CO_2_. Cell viability was determined by trypan blue exclusion before each experiment; samples with viability below 80% after overnight recovery were excluded.

### Patient PBMC cytokine-release assay

Rested PBMCs were plated at 2 x 10^5^ viable cells per well in 200 μL complete RPMI in round-bottom 96-well plates. Cells were preincubated for 30 minutes with vehicle control (0.1% DMSO), CA-23 (0.1, 0.3, or 1 μM), FR104 (1 μM), or abatacept (1 μM). T-cell activation was initiated with a submaximal concentration of soluble anti-human CD3 antibody (clone OKT3, 0.1 μg/mL). No exogenous anti-CD28 antibody was used.

After 24 hours, supernatants were collected, clarified by centrifugation, and stored at -80°C until analysis. IFN-γ, IL-2, TNF-α, IL-17A, and GM-CSF were quantified using a multiplex electrochemiluminescence assay per manufacturer’s protocol. Cytokine concentrations below the lower limit of quantification were assigned the lower limit of quantification for analysis. For each donor, cytokine responses were normalized to the matched anti-CD3-stimulated vehicle condition.

### PBMC flow-cytometric phenotyping

For flow-cytometric analyses, PBMCs were cultured as described above for 48 hours. Brefeldin A (5 μg/mL) and monensin (2 μM) were added during the final 5 hours. Cells were stained with Zombie Aqua fixable viability dye, followed by surface antibodies against CD3, CD4, CD8, CD69, CD25, and CD127. Cells were fixed and permeabilized using the Foxp3/Transcription Factor Staining Buffer Set and stained intracellularly for Ki-67, FOXP3, IFN-γ, and IL-17A.

Samples were acquired on a BD FACSymphony A5 flow cytometer with compensation determined using single-stained UltraComp eBeads. Data were analyzed with FlowJo version 11. Lymphocytes were identified by forward- and side-scatter properties, followed by exclusion of doublets and nonviable cells. Activated, proliferating, Th1, Th17, and double-producing cells were quantified as CD69, Ki-67, IFN-γ, IL-17A, and IFN-γIL-17A cells, respectively, within the CD4 T-cell gate.

### Human whole-blood agonism and antagonism assay

Fresh sodium-heparinized whole blood from healthy adult donors (from STEMCELL Technologies) was diluted 1:1 with complete RPMI and plated at 250 μL per well in 96-well plates. For assessment of independent agonist activity, diluted blood was incubated with vehicle or CA-23 at 0.1, 0.3, and 1 μM without added stimulus. For assessment of CD28-dependent antagonism, blood was pretreated with vehicle or compound for 30 minutes and stimulated with anti-CD3 antibody (0.1 μg/mL) together with recombinant human CD80-Fc and CD86-Fc (1 μg/mL each).

After 24 hours, supernatants were analyzed for IFN-γ, IL-2, and TNF-α. For CD69 analysis, cells were incubated for 18 hours, erythrocytes were lysed with ammonium chloride-based red blood cell lysis buffer, and leukocytes were stained for viability, CD3, CD4, CD8, and CD69. A compound was classified as lacking independent agonist activity when cytokine release and CD69 expression did not differ from unstimulated vehicle-treated blood.

### Autologous Treg suppression assay

CD4^+^ T cells were enriched from rested PBMCs by negative magnetic selection. Tregs were purified by fluorescence-activated cell sorting. Autologous responder T cells were sorted and labeled with CellTrace Violet. CD14^+^ monocytes were isolated by magnetic selection and used as autologous antigen-presenting cells.

Responder T cells, Tregs, and monocytes were cultured at a 2:1:1 ratio in complete RPMI. Cultures were pretreated for 30 minutes with vehicle, CA-23 (1 μM), FR104 (1 μM), or abatacept (1 μM), then stimulated with anti-CD3 antibody (0.1 μg/mL). Responder-only wells were included for each treatment condition to calculate Treg-mediated suppression. After 96 hours, CellTrace dilution was measured. Each condition was tested in duplicate, and duplicate values were averaged before donor-level analysis.

### Human intestinal epithelial-PBMC Transwell co-culture

Ready-to-use primary human small-intestinal epithelial tissues (EpiIntestinal^TM^ SMI-100; MatTek Life Sciences) were transferred to 24-well culture plates containing maintenance medium and equilibrated per manufacturer’s instructions before initiation of epithelial–PBMC co-cultures. Monolayers were equilibrated in complete epithelial maintenance medium, and only inserts with stable pretreatment TEER values were included. Rested PBMCs were added to the basolateral compartment at 5 × 10^5^ cells per insert in 500 μL complete medium. Vehicle, CA-23 (1 μM), FR104 (1 μM), or abatacept (1 μM) was added to the basolateral compartment 30 minutes before anti-CD3 stimulation (0.1 μg/mL).

TEER was measured immediately before treatment and after 48 hours using an EVOM3 voltohmmeter with STX2 electrodes. Resistance values from cell-free inserts were subtracted, and values were normalized to the corresponding pretreatment TEER for each epithelial insert. For permeability measurements, 4-kDa FITC-dextran was added to the apical chamber at 1 mg/mL after the 48-hour co-culture period. Basolateral medium was collected 2 hours later, and fluorescence was measured at 485/535 nm. FITC-dextran permeability was expressed relative to the anti-CD3-stimulated vehicle condition.

### Autologous monocyte-T-cell co-culture

CD14^+^ monocytes and CD4^+^ T cells were isolated from the same PBMC donor by magnetic selection. Monocytes were stimulated with lipopolysaccharide (100 ng/mL) for 18 hours to induce antigen-presenting-cell activation, washed twice, and cocultured with autologous CD4^+^ T cells at a 1:1 ratio. Vehicle or CA-23 at 0.1, 0.3, or 1 μM was added 30 minutes before stimulation with anti-CD3 antibody (0.1 μg/mL).

After 24 hours, CD4^+^ T-cell activation was assessed by CD69 expression and IL-2 secretion. Monocytes were stained for CD14, CD80, and CD86. CD80/CD86 expression was reported as median fluorescence intensity (MFI) after subtraction of matched fluorescence-minus-one controls.

### Surface plasmon resonance

CA-23 binding to immobilized human CD28, mouse CD28, and human CTLA-4 was measured by SPR (Biacore T200). Full instrument parameters are provided in the Supplementary Materials and Methods.

### In vitro ADME and early safety studies

In vitro ADME properties (kinetic solubility, PAMPA permeability, chemical and metabolic stability, plasma protein binding) and early safety parameters (WI-38/HS-27 cytotoxicity, hERG inhibition) were evaluated using standard protocols detailed in the Supplementary Materials and Methods; full results are summarized in Table S3.

### Animal studies

All animal experiments were approved by the Institutional Animal Care and Use Committee (IACUC) of our institution (protocol no. 2023-0028) and conducted in accordance with the institution’s animal welfare guidelines. C57BL/6J donor mice and C57BL/6J Rag1^-/-^ recipient mice were purchased from The Jackson Laboratory (Bar Harbor, ME, USA) and housed under specific pathogen-free conditions with a 12-h light/dark cycle and access to food and water.

### Single-dose IP pharmacokinetics and tissue distribution

Eight- to ten-week-old C57BL/6J mice received a single IP administration of CA-23 at 10 mg/kg in PBS containing 5% DMSO. Mice were randomly assigned to terminal sampling time points, with three mice per time point and balanced representation of both sexes. Blood was collected at predose and 0.083, 0.25, 0.5, 1, 2, 4, 8, and 24 hours after dosing. Colon and mesenteric lymph nodes were collected at 0.5, 2, 6, and 24 hours.

Whole blood was collected into K-EDTA tubes, centrifuged at 2,000 × g for 10 minutes at 4°C, and plasma was stored at -80°C. Colon and mesenteric lymph-node samples were rinsed in ice-cold PBS, weighed, snap-frozen, and stored at -80°C. Tissues were homogenized in four volumes of ice-cold 50% methanol/water. Plasma and tissue homogenates were spiked with internal standard, extracted by protein precipitation with four volumes of acetonitrile, centrifuged at 16,000 × g for 10 minutes, and analyzed by LC-MS/MS.

Calibration standards and quality-control samples were prepared in matched blank plasma, colon homogenate, and MLN homogenate. Quantification was based on the analyte-to-internal-standard peak-area ratio using weighted linear regression. Noncompartmental pharmacokinetic parameters, including AUC, and terminal half-life, were calculated using Phoenix WinNonlin. Tissue concentrations were reported as nmol/g, and tissue-to-plasma exposure ratios were calculated using AUC values.

### CD4^+^CD45RB^hi^ T cell adoptive transfer model of colitis

Chronic colitis was induced by adoptive transfer of CD4 positive and CD45RB high population (CD4+CD45RBhi) T cells into Rag1-/- recipient mice, essentially as described previously (*31–33*). Briefly, spleens were harvested from 5-week-old C57BL/6J donor mice. Single-cell suspensions were prepared by mechanical dissociation followed by filtration through a 70-μm nylon mesh. CD4^+^ T cells were enriched by magnetic-bead negative selection using Dynabeads™ Untouched™ Mouse CD4 Cells Kit (Invitrogen), and CD4^+^CD45RBhi T cells were further purified by fluorescence-activated cell sorting (FACS) on a Symphony S6 SE cell sorter using anti-CD4 (clone GK1.5) and anti-CD45RB (clone C363-16A) antibodies; the CD45RBhi gate was defined as the top 10-15% of CD45RB expressing CD4^+^ T cells. Sorted cells were washed twice in sterile PBS and resuspended at 2 × 10^6^ cells/ml. Each 6-7 week old Rag1⁻/⁻ recipient mouse received 4 × 10^5^ CD4^+^CD45RBhi T cells intraperitoneally (200 μl total volume). After cell transfer, mice were randomly assigned to treatment groups (n = 5-7 per group). CA-23 was dissolved in sterile PBS containing 5% DMSO and administered daily by intraperitoneal injection at doses of 1 or 10 mg/kg. Vehicle-treated mice received the same volume of formulation buffer (PBS containing 5% DMSO). Negative control mice received daily vehicle injections on the same schedule but did not undergo T cell transfer. Treatment began on day 1 post-transfer and continued daily for 45 days. Body weight was recorded every 3 days and expressed as a percentage of day 0 weight. At study termination (day 45), mice were euthanized, and colons were excised from cecum to rectum and measured under standardized tension. A portion of each colon was prepared as a Swiss roll, fixed in 4% paraformaldehyde, paraffin-embedded, sectioned at 5 μm, and stained with hematoxylin and eosin (H&E). Histological scoring was performed in a blinded manner using a modified version of the scoring system described by Erben et al. as follows: 0 = normal; 1 = minimal hyperplasia; 2 = mild hyperplasia, minimal goblet cell loss ± erosions; 3 = moderate hyperplasia ± few crypt abscesses, moderate goblet cell loss ± erosions; 4 = marked hyperplasia ± several crypt abscesses and/or erosions; 5 = marked hyperplasia ± multiple crypt abscesses (*34*).

### Flow cytometry of spleen and mesenteric lymph node lymphocytes

At study termination, spleens and mesenteric lymph nodes (MLN) were harvested from each mouse. Single-cell suspensions were prepared by mechanical dissociation through a 70-μm nylon mesh, followed by red blood cell lysis (spleen only) using RBC lysis buffer (BioLegend). Cells were resuspended at 2 × 10⁶ cells/ml in complete RPMI 1640 medium and stimulated for 5 h at 37 °C with phorbol 12-myristate 13-acetate (PMA, 1 μg/ml) and ionomycin (50 ng/ml) in the presence of brefeldin A (BD GolgiPlug™, BD Biosciences) per manufacturer’s protocol. Stimulated cells were stained for viability with Zombie Aqua™ Fixable Viability Kit (BioLegend) for 15 min at room temperature, followed by surface staining with anti-CD4 (clone RM4-5, PerCP-eFluor 710, eBioscience/Invitrogen) for 30 min at 4 °C. Cells were then fixed and permeabilized using the BD Cytofix/Cytoperm™ Plus Fixation/ Permeabilization Kit (BD Biosciences) per manufacturer’s instructions, and stained intracellularly with anti-IL-17A (clone TC11-18H10.1, PE, BioLegend) and anti-IFN-γ (clone XMG1.2, APC, BD Pharmingen) for 30 min at 4 °C. Compensation was performed using UltraComp eBeads™ Plus Compensation Beads (Invitrogen). Data were acquired on a BD FACSymphony™ A5 Cell Analyzer (BD Biosciences) and analyzed using FlowJo software v11 (BD Biosciences).

### Statistical analysis

Each human donor and each mouse was treated as one independent biological replicate. Technical replicates were averaged before statistical testing. Results are presented as mean ± SD unless stated otherwise. Comparisons among multiple treatment groups were performed using one-way analysis of variance with Tukey’s multiple-comparisons test or two-way analysis of variance when appropriate. Longitudinal body-weight measurements were analyzed using a mixed-effects model with treatment and time as fixed effects and animal as a repeated factor. Concentration-response curves were fitted using four-parameter logistic regression. Pharmacokinetic parameters were calculated by noncompartmental analysis. All statistical tests were two-sided, and p < 0.05 was considered statistically significant.

## List of Supplementary Materials

Supplementary Materials and Methods

Fig. S1 to Fig. S9

Table S1 to Table S3

LC-MS and NMR spectra

## Acknowledgments

The authors thank members of the Gabr Laboratory for helpful discussions.

## Author contributions

Conceptualization: SC, MG

Methodology: SU, SY, SC

Investigation: SU, SY, SC

Formal analysis: SC

Data curation: SU, SY, SC

Writing - original draft

SC Writing - review & editing

MG Supervision: MG

Funding acquisition: MG

## Data and materials availability

All data needed to evaluate the conclusions in this paper are available in the main text or the supplementary materials.

