## Supporting Information for "A small molecule inhibitor of CD28 costimulation restrains pathogenic T-cell responses in inflammatory bowel disease"

#### Supplementary Materials and Methods

##### Compounds and reagents

The Discovery Diversity Set (4,960 compounds) was purchased from Enamine. Abatacept was obtained from MedChemExpress (Monmouth Junction, NJ, USA). Furimazine was obtained from AOBIOUS (Gloucester, MA, USA). All other compounds were obtained from Sigma-Aldrich (St. Louis, MO, USA), unless otherwise specified. Recombinant human CD28 was purchased from ACROBiosystems (Newark, DE, USA), and recombinant human CD80, CD86, and CTLA-4 proteins were purchased from Bio-Techne (Minneapolis, MN, USA).

##### Surface plasmon resonance

SPR measurements were performed using a Biacore T200 instrument at 25°C. Recombinant human CD28, mouse CD28, and human CTLA-4 extracellular domains were individually immobilized on CM5 sensor-chip flow cells by standard amine coupling. Proteins were diluted in 10 mM sodium acetate buffer, pH 5.0, and immobilized on the surface of the chip. A reference flow cell was activated and quenched without protein immobilization.

CA-23 was dissolved in running buffer consisting of HBS-EP containing 1% DMSO and injected at 0.078, 0.313, 1.25, 5, and 10 μM. Each injection included a 150-second association phase and 300-second dissociation phase at a flow rate of 30 μL/min. Blank-buffer injections were interspersed throughout each run, and a DMSO calibration series was used for solvent correction. Sensorgrams were double referenced by subtraction of the reference flow cell and blank-buffer responses.

Human and mouse CD28 binding data were fitted using a 1:1 Langmuir kinetic model when residuals were randomly distributed and values met prespecified quality criteria. CTLA-4 was considered nonbinding when reference-subtracted responses remained within assay noise and did not increase with CA-23 concentration.

**In vitro ADME and early safety studies**

Kinetic solubility of CA-23 was determined in 1% DMSO/PBS, pH 7.4, and fasted-state simulated intestinal fluid. Passive permeability was evaluated by PAMPA using a phospholipid-coated artificial membrane. Chemical stability was assessed in simulated gastric fluid, pH 1.2, and simulated intestinal fluid, pH 6.8, after 2 hours at 37°C. Metabolic stability was measured in pooled mouse and human liver microsomes in the presence of NADPH. Plasma stability was determined in pooled mouse and human plasma at 37°C. Compound concentrations were measured by LC-MS/MS, and half-lives were calculated by log-linear regression of the percentage of parent compound remaining over time.

Human plasma protein binding was determined by equilibrium dialysis. Cytotoxicity in WI-38 and HS-27 cells was measured after 72-hour compound exposure using a luminescent ATP-content assay. hERG inhibition was measured by automated whole-cell patch clamp in HEK293 cells stably expressing hERG channels. Concentration-response curves were fitted by four-parameter logistic regression.

**Table S1. Structure-activity relationship of CA-01 derivatives and their $IC_{50}$ values against CD28-CD80 and CD28-CD86 interactions.**

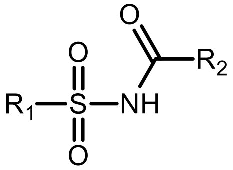


| Name | $R_1$ | $R_2$ | $IC_{50}$ (μM) | |
|---|---|---|---|---|
| | | | CD80 | CD86 |
| CA-01 | 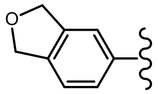 | 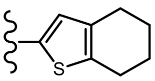 | 0.283 ± 0.03 | 0.547 ± 0.21 |
| CA-02 | 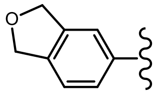 | 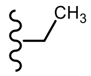 | > 10 | > 10 |
| CA-03 | 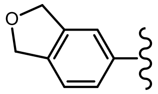 | 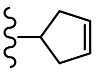 | > 10 | > 10 |
| CA-04 | 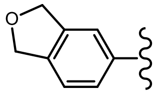 | 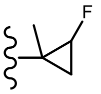 | > 10 | > 10 |
| CA-05 | 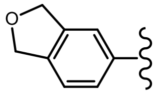 | 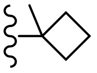 | > 10 | > 10 |
| CA-06 | 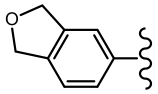 | 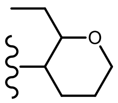 | > 10 | > 10 |
| CA-07 | 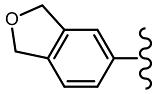 | 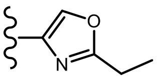 | > 10 | > 10 |
| CA-08 | 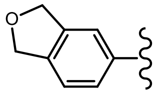 | 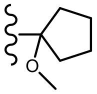 | > 10 | > 10 |

| | | | | |
|---|---|---|---|---|
| CA-09 | 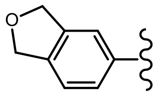 | 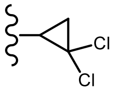 | > 10 | > 10 |
| CA-10 | 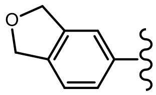 | 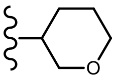 | > 10 | > 10 |
| CA-11 | 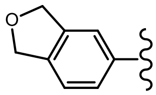 | 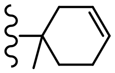 | > 10 | > 10 |
| CA-12 | 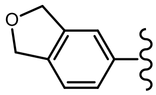 | 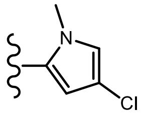 | ~ 10 | ~ 10 |
| CA-13 | 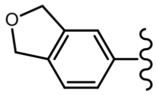 | 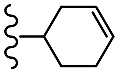 | > 10 | > 10 |
| CA-14 | 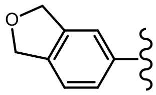 | 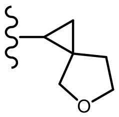 | > 10 | > 10 |
| CA-15 | 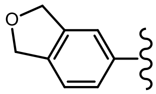 | 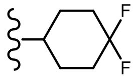 | > 10 | > 10 |
| CA-16 | 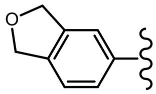 | 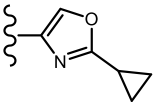 | > 10 | > 10 |
| CA-17 | 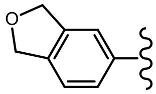 | 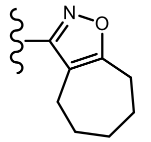 | > 10 | > 10 |
| CA-18 | 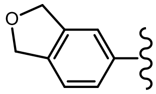 | 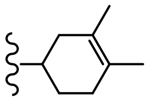 | ~ 10 | ~ 10 |
| CA-19 | 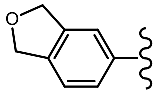 | 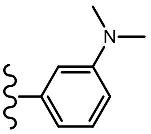 | > 10 | > 10 |

| | | | | |
|---|---|---|---|---|
| CA-20 | 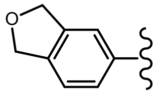 | 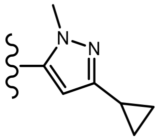 | > 10 | > 10 |
| CA-21 | 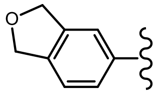 | 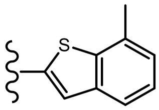 | 0.32 ± 0.19 | 0.24 ± 0.07 |
| CA-22 | 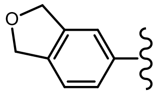 | 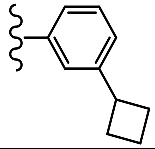 | 2.8 ± 1.93 | 3.45 ± 0.9 |
| CA-23 | 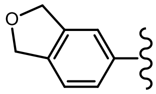 | 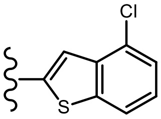 | 0.035 ± 0.004 | 0.024 ± 0.003 |
| CA-24 | 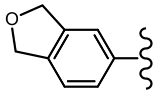 | 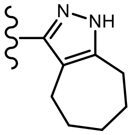 | > 10 | > 10 |
| CA-25 | 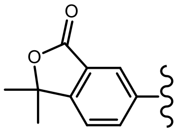 | 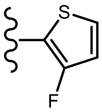 | > 10 | > 10 |
| CA-26 | 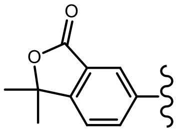 | 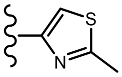 | > 10 | > 10 |
| CA-27 | 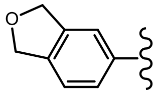 | 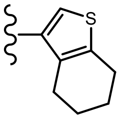 | 4.93 ± 2.17 | 1.56 ± 0.7 |
| CA-28 | 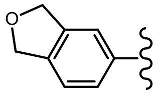 | 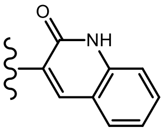 | > 10 | > 10 |
| CA-29 | 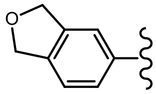 | 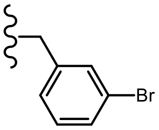 | > 10 | > 10 |
| CA-30 | 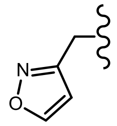 | 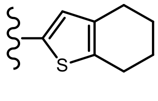 | > 10 | > 10 |

| | | | | |
|---|---|---|---|---|
| CA-31 | 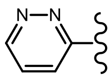 | 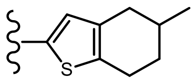 | > 10 | > 10 |
| CA-32 | 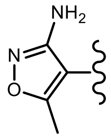 | 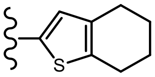 | ~ 10 | > 10 |
| CA-33 | 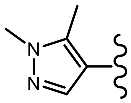 | 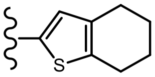 | 2.63 ± 0.36 | 2.84 ± 1.27 |

**Table S2. Structure-activity relationship of CB-01 derivatives and their $IC_{50}$ values against CD28-CD80 and CD28-CD86 interactions.**

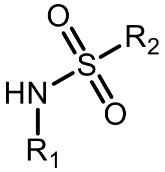


| Name | $R_1$ | $R_2$ | $IC_{50}$ (μM) | |
|---|---|---|---|---|
| | | | CD80 | CD86 |
| CB-01 | 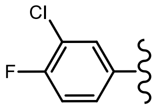 | 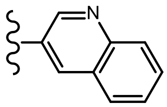 | 0.35 ± 0.16 | 0.4 ± 0.06 |
| CB-02 | H | 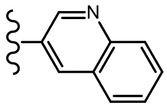 | > 10 | > 10 |
| CB-03 | 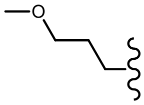 | 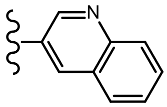 | > 10 | > 10 |
| CB-04 | 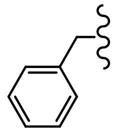 | 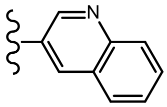 | > 10 | > 10 |
| CB-05 | 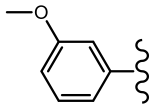 | 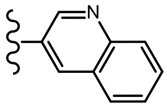 | > 10 | > 10 |
| CB-06 | 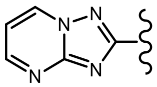 | 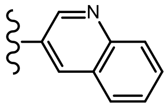 | > 10 | > 10 |
| CB-07 | 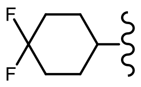 | 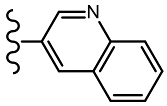 | > 10 | > 10 |
| CB-08 | 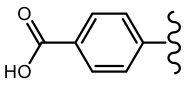 | 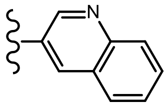 | > 10 | > 10 |
| CB-09 | 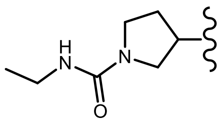 | 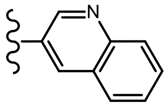 | > 10 | > 10 |

| | | | | |
|---|---|---|---|---|
| CB-10 | 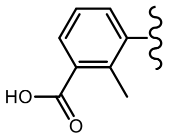 | 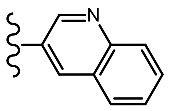 | > 10 | > 10 |
| CB-11 | 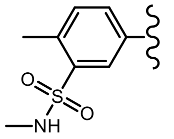 | 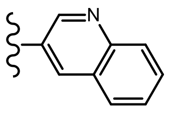 | > 10 | > 10 |
| CB-12 | 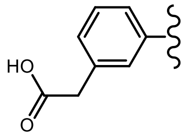 | 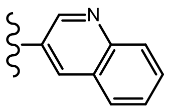 | > 10 | > 10 |
| CB-13 | 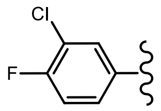 | 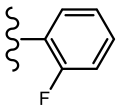 | ~ 10 | ~ 10 |
| CB-14 | 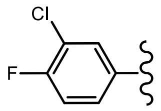 | 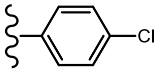 | 8.6 ± 3.93 | 5.37 ± 2.37 |
| CB-15 | 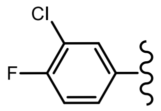 | 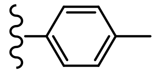 | ~ 10 | ~ 10 |
| CB-16 | 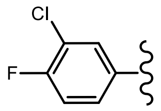 | 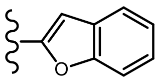 | 0.45 ± 0.15 | 0.52 ± 0.26 |
| CB-17 | 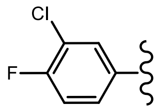 | 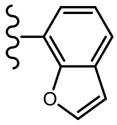 | > 10 | > 10 |
| CB-18 | 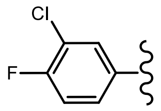 | 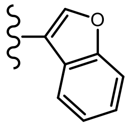 | 3.25 ± 1.27 | 1.38 ± 0.13 |
| CB-19 | 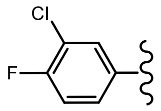 | 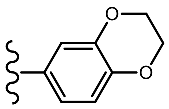 | 2.73 ± 1.45 | 1.05 ± 0.58 |
| CB-20 | 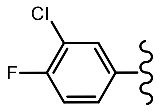 | 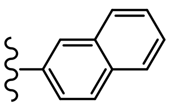 | 2.72 ± 1.28 | 1.75 ± 0.16 |

| | | | | |
|---|---|---|---|---|
| CB-21 | 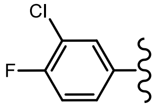 | 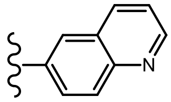 | 5.82 ± 2.64 | 2.39 ± 0.31 |
| CB-22 | 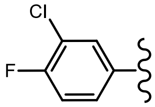 | 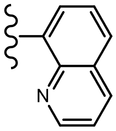 | > 10 | > 10 |
| CB-23 | 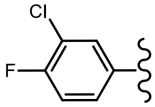 | 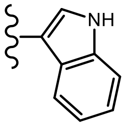 | ~ 10 | ~ 10 |
| CB-24 | 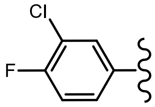 | 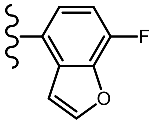 | 3.45 ± 1.36 | 2.52 ± 0.2 |
| CB-25 | 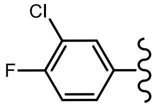 | 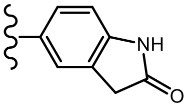 | > 10 | > 10 |
| CB-26 | 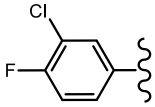 | 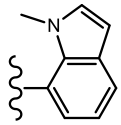 | 1.33 ± 0.76 | 1.02 ± 0.34 |
| CB-27 | 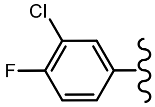 | 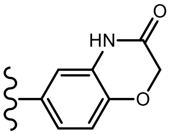 | ~ 10 | ~ 10 |
| CB-28 | 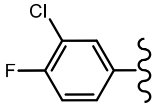 | 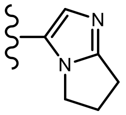 | > 10 | > 10 |
| CB-29 | 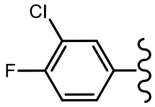 | 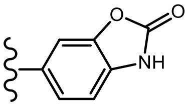 | 1.87 ± 0.23 | 2.53 ± 1.84 |
| CB-30 | 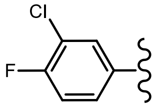 | 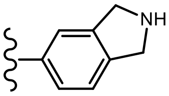 | > 10 | > 10 |
| CB-31 | 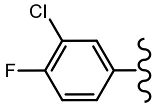 | 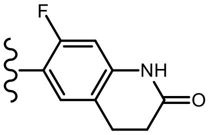 | ~ 10 | ~ 10 |

| | | | | |
|---|---|---|---|---|
| CB-32 | 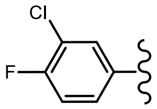 | 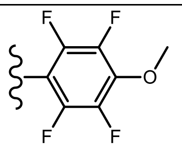 | 0.014 ± 0.003 | 0.005 ± 0.002 |
| CB-33 | 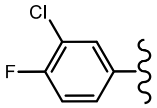 | 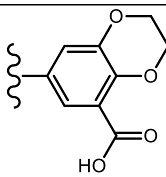 | > 10 | > 10 |

**Table S3. In vitro absorption, distribution, metabolism, excretion, and early safety profile of CA-23.**

| PK Parameter | CA-23 |
|---|---|
| $LogD_{7.4}$[a] | 3.1 |
| Solubility, 1% DMSO/PBS [µM][b] | 85 |
| Solubility in FaSSIF [µM] | 152 |
| PAMPA $P_{app}$ [$\times10^{-6}$ cm/s] | 12.6 |
| Stability in simulated gastric fluid (pH 1.2, 2 h), % remaining | 97.2 |
| Stability in simulated intestinal fluid (pH 6.8, 2 h), % remaining | 98.1 |
| t½ [min], mouse liver microsomes | 68.0 |
| t½ [min], human liver microsomes | 84.0 |
| t½ [min], mouse plasma | >100[c] |
| t½ [min], human plasma | >100[c] |
| t½ [min], PBS | >100[c] |
| Human plasma protein binding [%] | 94.0 |
| $IC_{50}$ against WI-38 [µM] | >100 |
| $IC_{50}$ against HS-27 [µM] | >100 |
| $IC_{50}$ against hERG [µM] | 98.3 |

[a] LogD at physiological pH. [b] Kinetic-solubility format. [c] The half-life was not reached within a 100-min assay window.

### Supplementary Figures

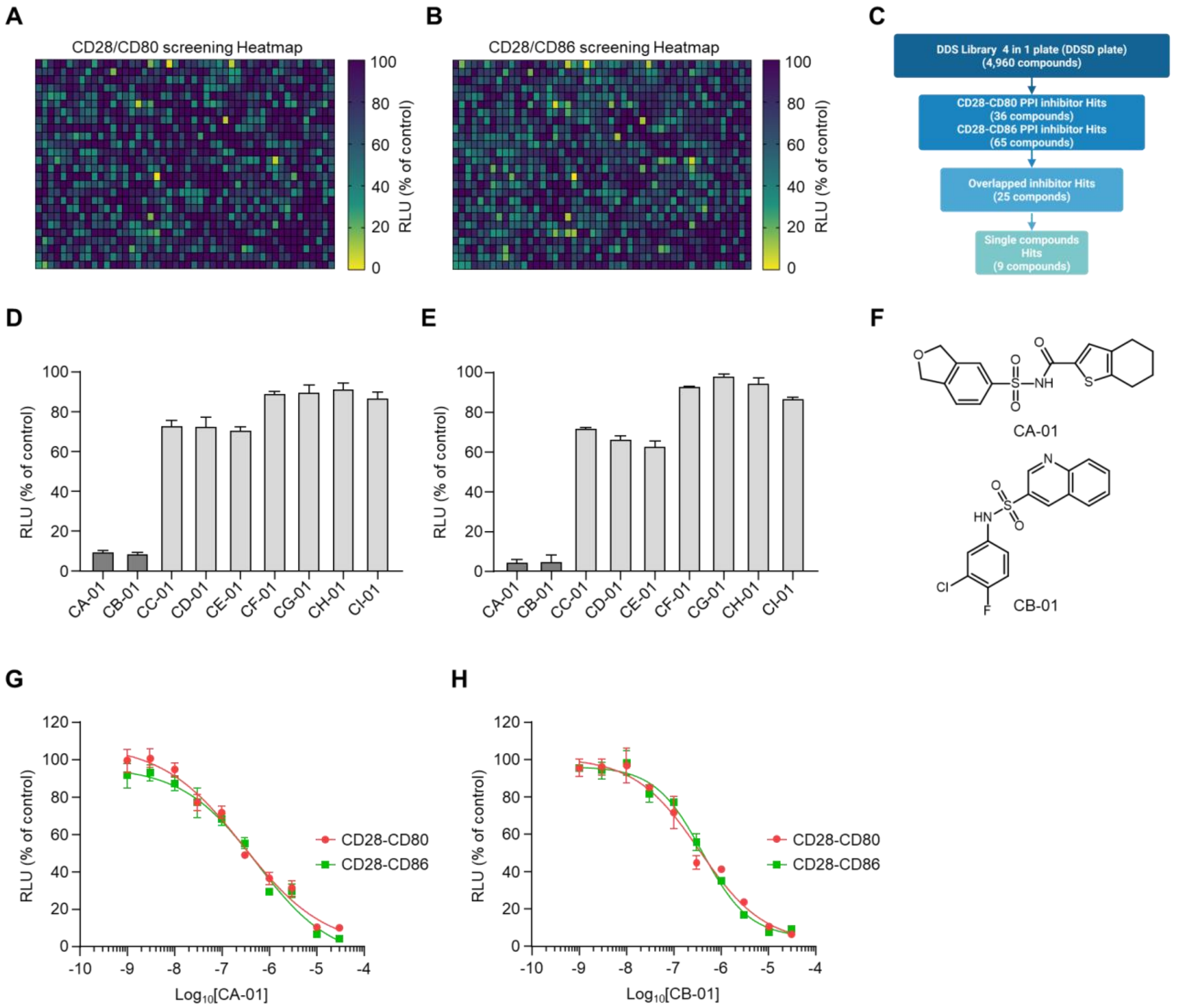


**Supplementary Figure 1. High-throughput screening identifies small molecule inhibitors of CD28-CD80 and CD28-CD86 interactions.**

(A, B) Heatmap of primary screening results for CD28-CD80 (A) and CD28-CD86 (B) interactions using the NanoBiT assay. Compounds were screened at 25 μM for 3 hours. Luminescence is shown as % of vehicle control (DMSO).

(C) Hit selection flowchart. Hits were defined as compounds reducing luminescence to <30% of vehicle control.

(D, E) Confirmation of 9 single-compound hits against CD28-CD80 (D) and CD28-CD86 (E) interactions at 10 μM. Data are presented as mean ± SD (n=3).

(F) Chemical structures of the two hit compounds, CA-01 and CB-01.

(G, H) Dose-response curves of CA-01 (G) and CB-01 (H) against CD28-CD80 (red) and CD28-CD86 (green) interactions. Data are presented as mean ± SD (n=5 or more).

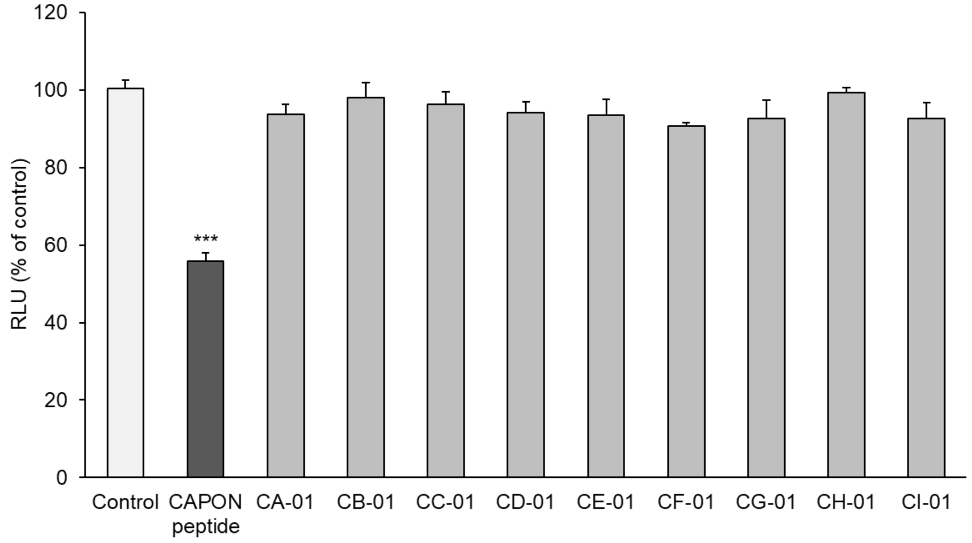


**Supplementary Figure 2. Selectivity of CD28-B7 PPI inhibitor hits against an unrelated NanoBiT-based protein-protein interaction.**

CHO-K1 cells stably expressing NOS1-HiBiT and NOS1AP-LgBiT were treated with the indicated hit candidates (CA-01 to CI-01) at 100 μM, and luminescence was measured to evaluate inhibitory activity against the unrelated NOS1-NOS1AP interaction. TAT-GESV (10 μM), a known peptide inhibitor of the NOS1-NOS1AP interaction, was included as a positive control. Luminescence is shown as % of vehicle (DMSO) control. Data are presented as mean ± SD (n=3). ***$p < 0.001$ vs Control (one-way ANOVA with Dunnett's post-hoc test).

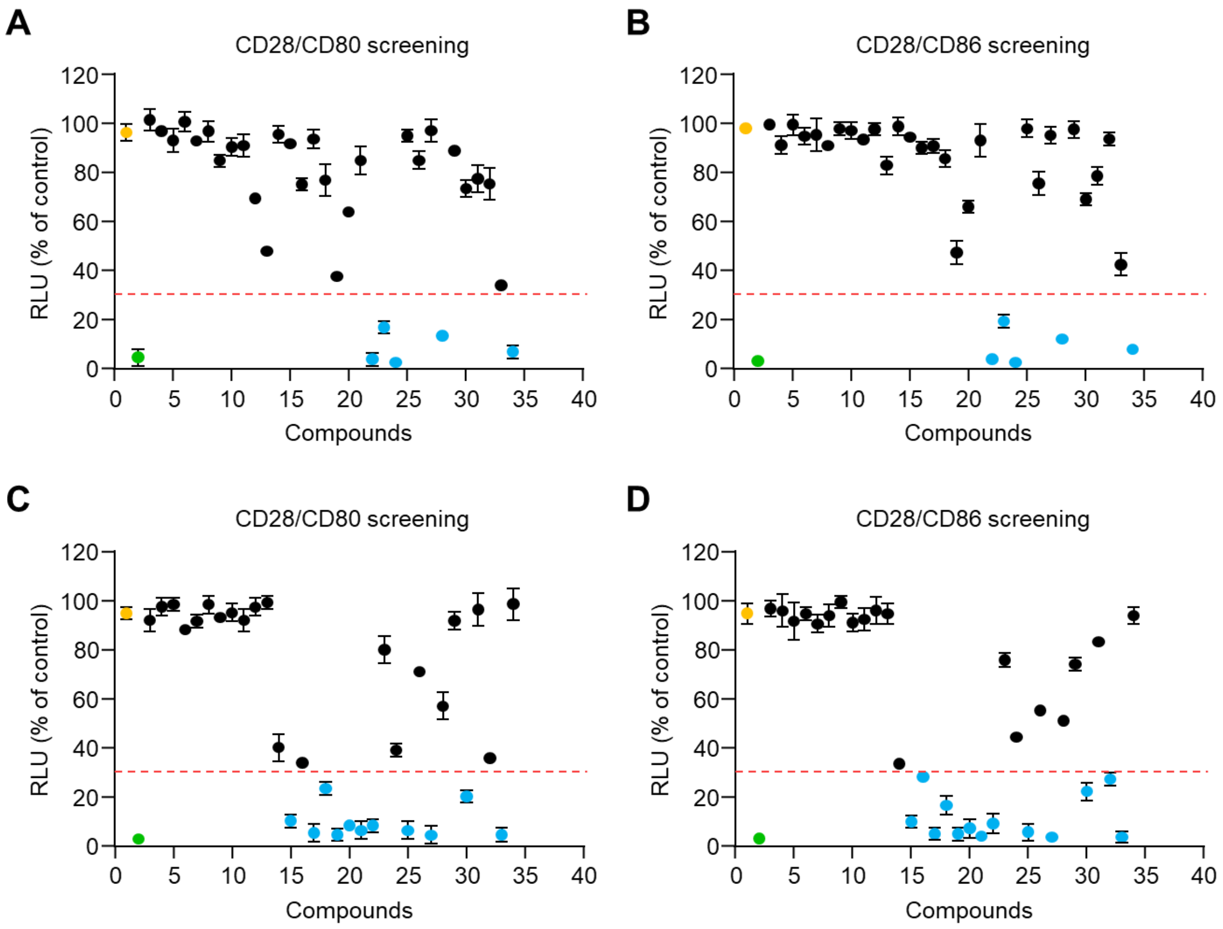


**Supplementary Figure 3. Structure-activity relationship (SAR) screening of CA-01 and CB-01 derivatives against CD28-CD80 and CD28-CD86 interactions.**

(A, B) Inhibitory activity of CA-01 32 derivatives against CD28-CD80 (A) and CD28-CD86 (B) interactions. Each dot represents an individual analog tested at 10 µM. The red dashed line indicates the hit threshold (30% of vehicle control). Yellow dots, vehicle (DMSO); green dots, parental CA-01; blue dots, analogs meeting the hit threshold.

(C, D) Inhibitory activity of CB-01 32 derivatives against CD28-CD80 (C) and CD28-CD86 (D) interactions, displayed as in (A, B). Orange dots, vehicle; green dots, parental CB-01; blue dots, analogs meeting the hit threshold. Data are presented as mean ± SD (n=3).

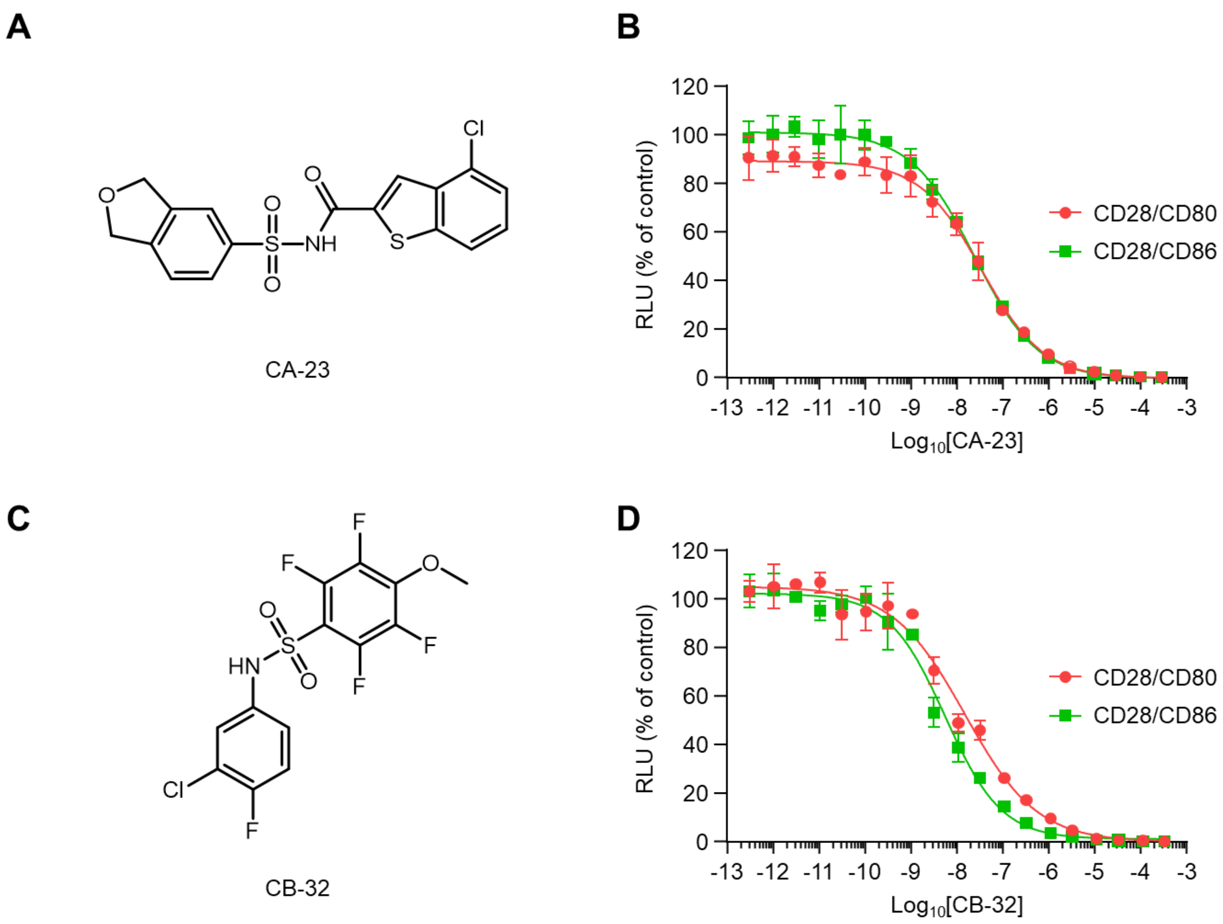


**Supplementary Figure 4. CA-23 and CB-32 are the most potent CD28-B7 PPI inhibitors derived from SAR optimization.**

(A, B) Chemical structure (A) and dose-response curves (B) of CA-23, the most potent compounds from the CA-01 derivatives.

(C, D) Chemical structure (C) and dose-response curves (D) of CB-32, the most potent compound from the CB-01 derivatives, Inhibition of CD28-CD80 (red line) and CD28-CD86 (green line) interactions was measured by the NanoBiT assay. Data are presented as mean ± SD (n=5)

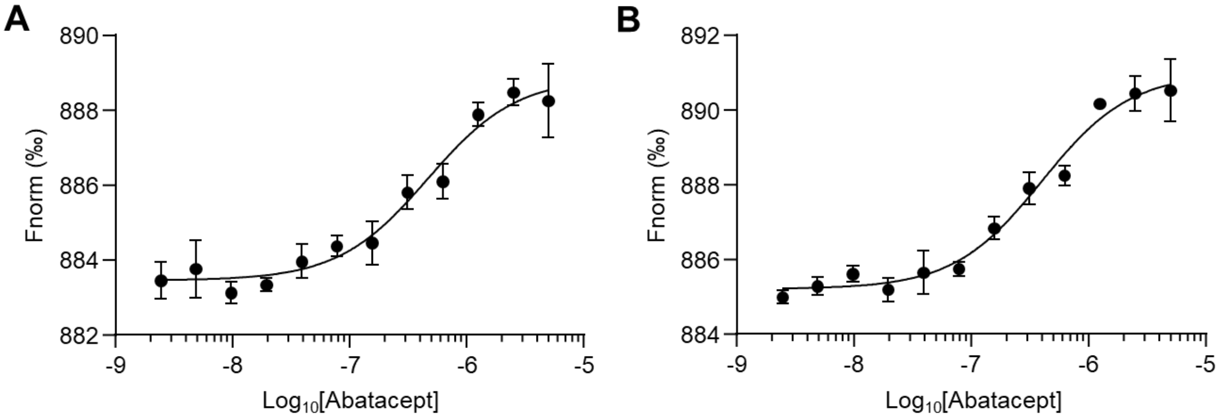


**Supplementary Figure 5. Abatacept binds CD80 and CD86 in a dose-dependent manner.**

(A, B) Microscale Thermophoresis (MST) dose-response curves of Abatacept binding to recombinant CD80 (A) and CD86 (B) proteins. Data are presented as mean ± SD (n=3).

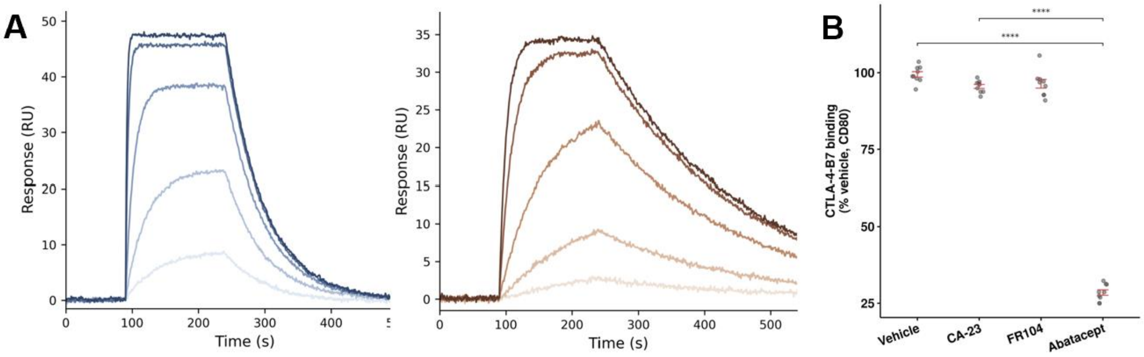


**Supplementary Figure 6. Species cross-reactivity of CA-23 by surface plasmon resonance (SPR) and functional CTLA-4-B7 competition.**

**(A)** SPR sensorgrams of CA-23 binding to immobilized human CD28 (left) and mouse CD28 (right) across a concentration series (0.078-10 μM), yielding $K_D$ = 0.35 μM (human) and 0.52 μM (mouse). **(B)** Functional CTLA-4-B7 competition assay: CD80 binding to CTLA-4 in the presence of vehicle, CA-23, FR104, or Abatacept (1 μM). Data are mean ± SEM with individual replicates shown; ****$p$<0.0001, t-test vs. vehicle.

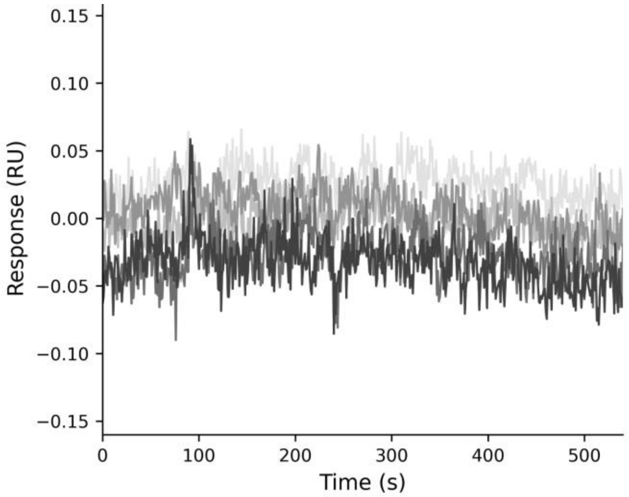


**Supplementary Figure 7. Human CTLA-4 shows no detectable binding to CA-23 by SPR.**

SPR sensorgrams of CA-23 across a concentration series (0.078-10 μM) against immobilized human CTLA-4, showing no dose-dependent response above assay noise (±1 RU).

**Supplementary Figure 8. Synthesis and characterization of CA-23**

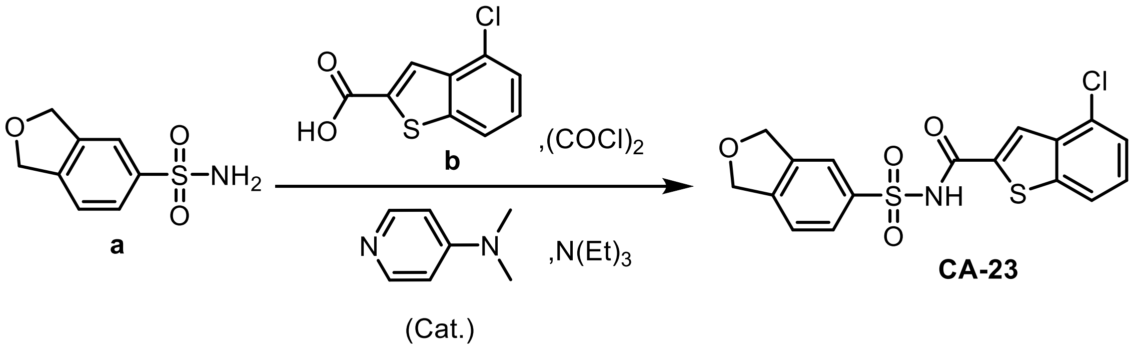


A 100 mL oven-dried round-bottom flask equipped with a stir bar was purged with dry $N_2$. 4-chlorobenzo[b]thiophene-2-carboxylic acid **b** (1.28 g, 6.02 mmol, 1.2 eq.), 2 drops of DMF and anhydrous DCM were added to the flask to form a suspension, followed by cooling to 0°C. Oxalyl chloride (0.828 g, 0.56 mL, 6.53 mmol, 1.3 eq.) was added dropwise to the flask over 5 minutes at 0°C. After complete addition, the reaction was maintained at 0°C until the reaction solution became completely clear. The mixture was maintained for additional one hour at 0°C and the solvent was removed under low pressure, followed by adding 10 mL anhydrous THF to dissolve the resulting solids. In a separate vial, 1,3-dihydroisobenzofuran-5-sulfonamide **a** (1.00 g, 5.02 mmol, 1 eq.) and DMAP (31 mg, 0.251 mmol, 5 mol%) were dissolved in 10 mL anhydrous THF, the resulting solution was slowly transferred to the acyl chloride solution at 0°C, followed by slowly adding triethylamine (0.66 g, 0.91 mL, 6.53 mmol, 1.3 eq.). The reaction was gradually warmed up to room temperature and kept for overnight. After the completion of the reaction, the mixture was concentrated under low pressure, redissolved in 200 mL 30% IPA/DCM, washed with 0.1M of HCl, saturated sodium bicarbonate solution and brine. The organic layer was dried over sodium sulfate, concentrated and purified under regular silica chromatography with 0%-80% hexanes/acetone to produce **CA-23** (1.25 g, 3.17 mmol, 63%) as light-yellow solid. $^{1}$**H NMR** (500 MHz, DMSO) δ 13.01 (s, 1H, *br*), 8.58 (s, 1H), 8.03 (d, $J$ = 8.0 Hz, 1H), 7.97 – 7.91 (m, 2H), 7.58 – 7.55 (m, 2H), 7.51 (t, $J$ = 7.9 Hz, 1H), 5.07 (dd, $J$ = 5.6, 2.1 Hz, 4H). $^{13}$**C NMR** (126 MHz, DMSO) δ 160.79, 145.78, 142.86, 140.77, 139.25, 138.75, 137.38, 129.72, 128.81, 127.76, 127.07, 125.63, 122.62, 122.38, 121.34, 72.75.

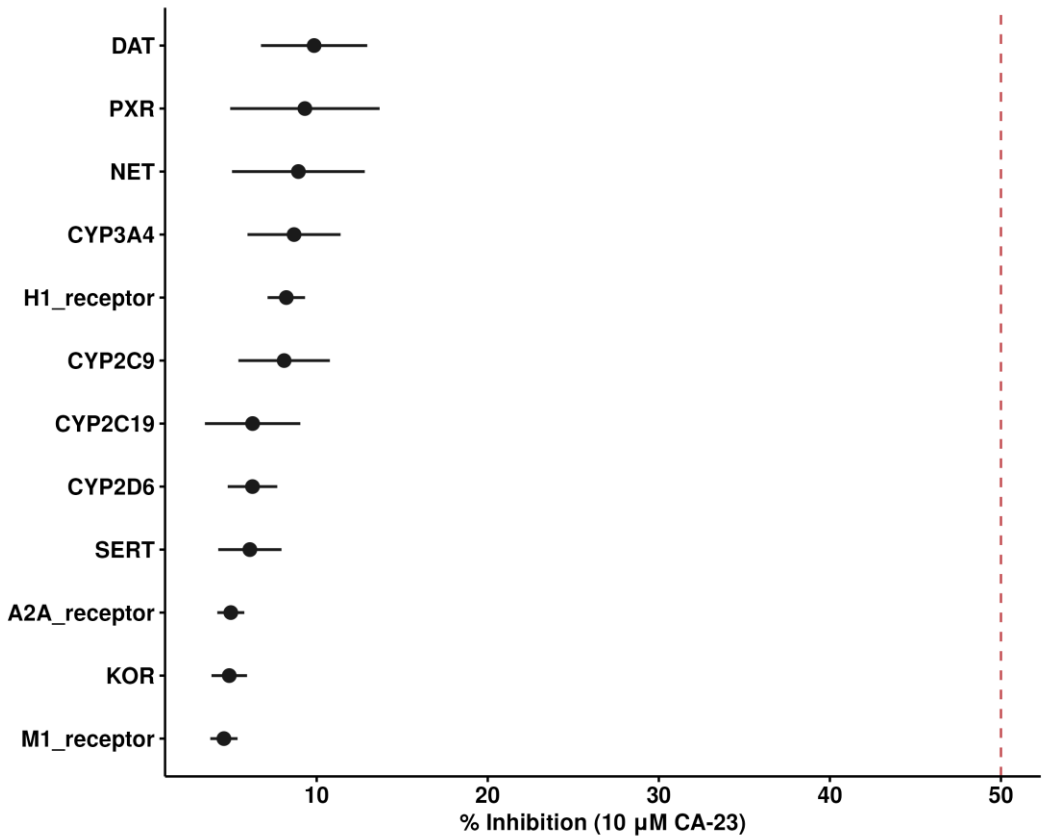


**Supplementary Figure 9. Off-target and CYP450 selectivity panel of CA-23**

Percent inhibition of CA-23 (10 μM) across a panel of CYP450 enzymes, GPCRs, and transporters (mean ± SD, n=3). Dashed line indicates the conventional 50% flag threshold; all targets showed <15% inhibition, indicating a >25-fold selectivity margin over the sub-micromolar potency of CA-23 at its intended target.

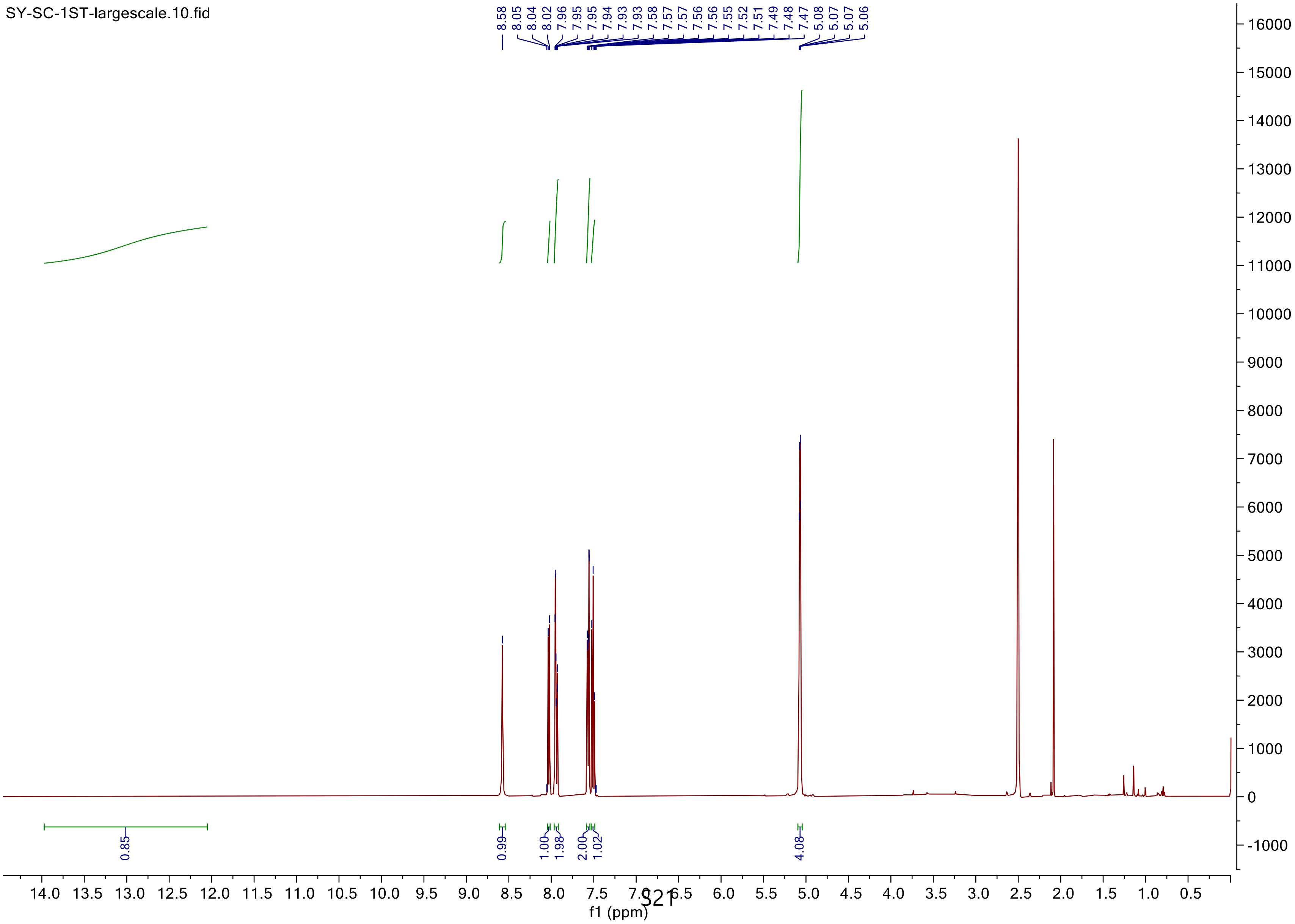
SY-SC-1ST-largescale.10.fid
8.58
8.05
8.04
8.02
7.96
7.95
7.95
7.94
7.93
7.93
7.58
7.57
7.57
7.56
7.56
7.55
7.52
7.51
7.49
7.48
7.47
5.08
5.07
5.07
5.06
0.85
0.99
1.00
1.98
2.00
1.02
4.08
f1 (ppm)

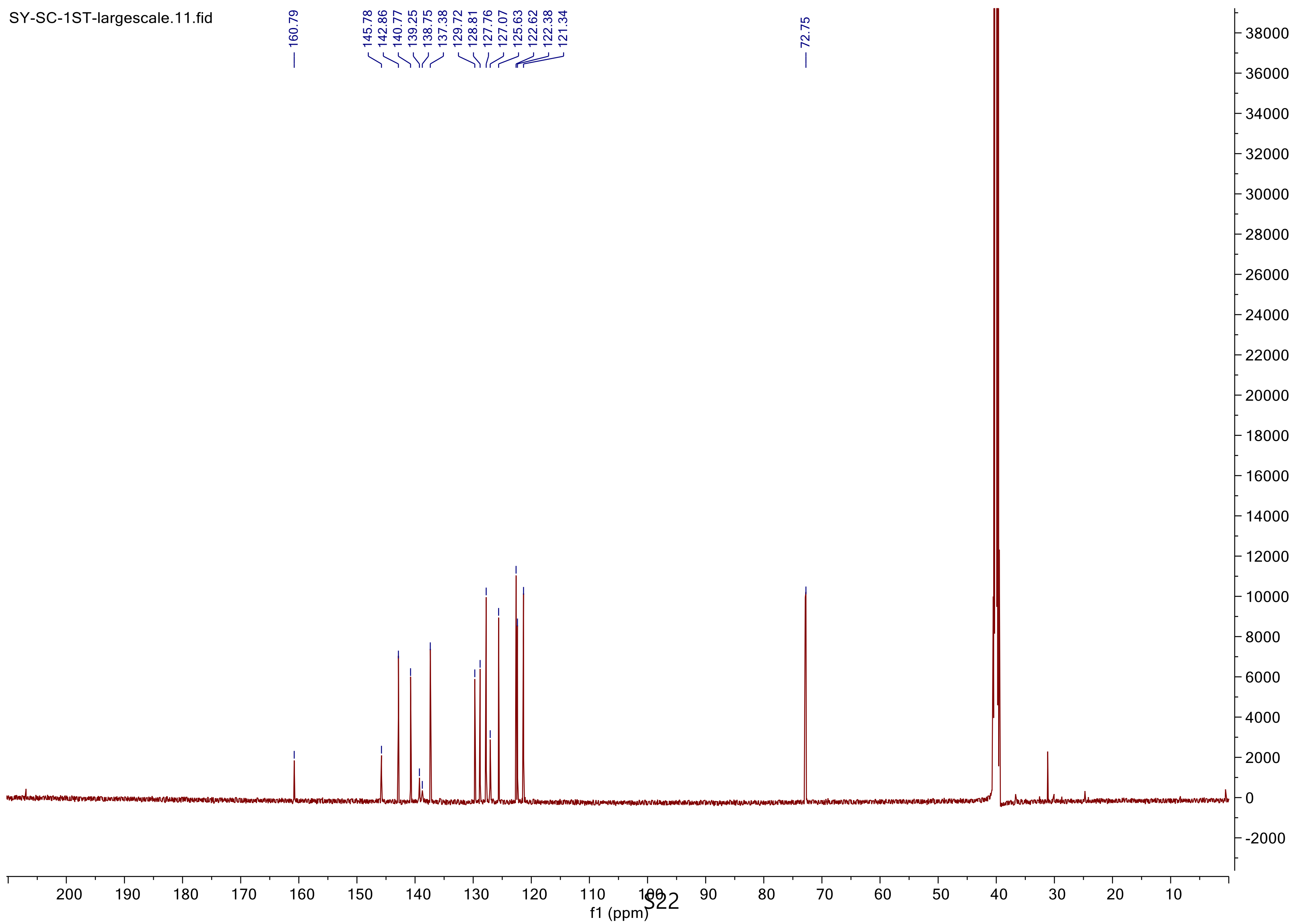
SY-SC-1ST-largescale.11.fid
160.79
145.78
142.86
140.77
139.25
138.75
137.38
129.72
128.81
127.76
127.07
125.63
122.62
122.38
121.34
72.75
f1 (ppm)

**LC-MS of CA-23**

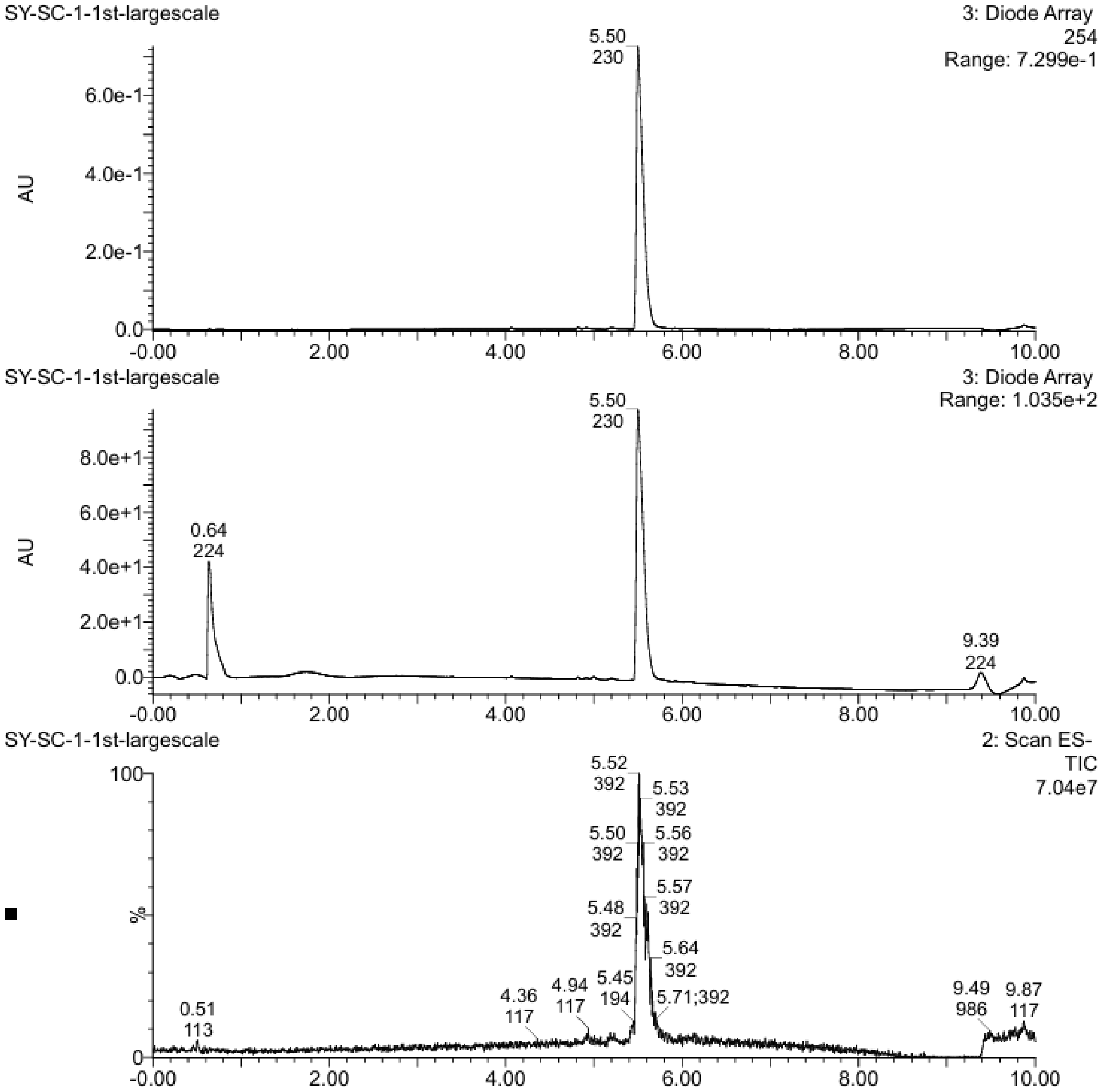
SY-SC-1-1st-largescale
3: Diode Array
254
Range: 7.299e-1
5.50
230
AU
6.0e-1
4.0e-1
2.0e-1
0.0
-0.00
2.00
4.00
6.00
8.00
10.00
SY-SC-1-1st-largescale
3: Diode Array
Range: 1.035e+2
5.50
230
0.64
224
9.39
224
8.0e+1
6.0e+1
4.0e+1
2.0e+1
0.0
SY-SC-1-1st-largescale
2: Scan ES-
TIC
7.04e7
100
%
0
5.52
392
5.53
392
5.50
392
5.56
392
5.57
392
5.48
392
5.64
392
5.45
194
5.71;392
4.36
117
4.94
117
0.51
113
9.49
986
9.87
117
